# Cis-inhibition suppresses basal Notch signalling during sensory organ precursor selection

**DOI:** 10.1101/2022.07.19.500683

**Authors:** Tobias Troost, Udi Binshtok, David Sprinzak, Thomas Klein

## Abstract

The emergence of the sensory organ precursor (SOP) from the proneural equivalence group in *Drosophila melanogaster* is a paradigm for studying single cell fate specification through the process of lateral inhibition. Classical lateral inhibition models describing this selection process are based on a transcriptional feedback mechanism where inhibitory signals between neighbouring cells, mediated by Notch pathway, are coupled to an intracellular circuit regulating the expression of the Notch ligand Delta (Dl). It was previously shown that in addition to its ability to trans-activate Notch in neighbouring cells, Dl can also cis-inhibit Notch in the same cell. However, it remains unclear what role does cis-inhibition play during SOP selection, and how it contributes to the selection of only one SOP. Here we address these questions using the unexpected observation that the mammalian ligand Delta-like 1 (Dll1) can trans-activate but not cis-inhibit Notch in *Drosophila*. We develop a mathematical model for SOP selection, termed the two-channel SOP (TCS) model, where Dl activity, but not its expression, is regulated by two channels associated with the two E3 ubiquitin ligases Neuralized (Neur) and Mindbomb1 (Mib1). While the Neur-dependent channel is regulated by Notch signalling, the Mib1-dependent channel is not, leading to tissue-wide basal inhibitory activity. We show theoretically and experimentally that cis-inhibition is required for suppressing Mib1-dependent basal Notch activity. Thus, our results highlight the trade-off between basal Notch activity and cis-inhibition as a mechanism for singling out an SOP from the proneural equivalence group.

## Introduction

The Notch signalling pathway is conserved across all metazoans and has been linked to numerous developmental, homoeostatic, and disease related processes (reviewed in [1]). In *Drosophila*, signalling is initiated by binding of one of its two Notch ligands, Delta (Dl) or Serrate (Ser), to the Notch receptor. Dl and Ser are type I transmembrane proteins, whose extracellular domains (ECD) consist of an N-terminal Delta/Serrate/Lag2 (DSL) domain, which mediates the interaction with Notch, followed by several repeats of the EGF motif and the transmembrane region. The binding of the ligands triggers an unusual transduction mechanism in which the Notch intracellular domain (NICD) is released into the cytosol, from which it is transported into the nucleus to become an essential component of a transcriptional activator complex that assembles around the DNA-binding factor Suppressor of Hairless (Su(H)), a member of the CSL transcription factor family.

Endocytosis of the ligand is required for the full activation of the pathway. Two E3 ligases have been identified, termed Mindbomb1 (Mib1) and Neuralized (Neur), which initiate endocytosis by ubiquitylating lysins (Ks) in the intracellular domain (ICD) of the ligands. *Cis*-inhibition (CI) is a mechanism that regulates the activity of the Notch pathway in a post-translational and concentration dependent manner whereby Notch ligands bind and inhibit Notch receptors in the same cell ([2–4], reviewed in [5]). It was first discovered in *Drosophila* in over-expression experiments and subsequently shown to act during several developmental processes to regulate the activity of the pathway in the wing, eye, oogenesis, and the notum in Drosophila (reviewed in [6]). Hence, CI is an important regulatory feature of Notch signalling. The mechanism of CI is not understood, but the DSL region of the ECD is essential [7, 8]. In Ser, the EGF repeats 4-6 of its ECD are required for CI, and the deletion of one of these repeats abolishes its cis-inhibitory abilities [9].

A classical, extensively studied, patterning process mediated by Notch signalling is the selection of sensory organ precursor cells (SOP) of the sensory bristle out of an equivalence group termed the proneural cluster (PNC). The PNCs in the notum of the fly are defined by the expression of the proneural genes *achaete* (*ac*) and *scute* (*sc*), which encode bHLH transcription factors (reviewed in [10]). Their expression imposes an undecided proneural state to the cells from which they can progress to develop into SOPs, or regress to the default epidermoblast state. Within a given PNC, only a defined number of cells (1 or 2) eventually adopt the SOP fate. They are selected by the activity of the Notch pathway in a process termed lateral inhibition. In the case of loss of Notch activity, all cells of a PNC adopt the SOP fate (neurogenic phenotype), indicating that Notch activity prevents the SOP fate in PNC cells by antagonising the activity of the proneural genes. According to the prevailing classical model of lateral inhibition the selection of SOP occurs through an intercellular transcriptional feedback mechanism, termed here the transcriptional feedback (TFB) model (reviewed in [11]). In the TFB model, Ac and Sc activate the expression of Dl, which activates the Notch pathway in its surrounding cells. Notch activity in each cell suppresses the expression of proneural genes and, hence, also of Dl. This feedback mechanism can amplify small initial differences in Ac and Sc expression among the PNC cells, such that one cell (the future SOP) starts expressing high Dl levels while its neighbours are suppressed. This regulatory feedback loop generates an all or none switch where one cell expresses high levels of proneural activity and becomes the SOP, whereas its neighbours switch to the epidermoblast fate.

Recent work provided strong evidence against the TFB model. Using the MARCM clone technique, a *Dl Ser* double mutant PNCs that also expressed either Dl or Ser at a uniform level in all mutant cells was generated [12]. Surprisingly, the selection of the SOP proceeded normally, although the transcription of Dl is uncoupled from the proneural activity. A similar observation has been previously made for the selection of the neuroblasts in the embryo [13]. These data suggest that differential expression of Dl, a crucial element in the classical TFB model, is dispensable for the SOP selection process. In line with this conclusion is the observation that the overall expression of Dl appears to be unchanged in *ac sc* mutants, indicating that the influence of the proneural factors on the expression of Dl is minimal [14, 15]. Moreover, the TFB model cannot explain how the SOP via a short-range signalling pathway (Notch) inhibit cells in the PNC that are not in direct contact with it. The TFB model can also not explain the difference in the loss of function phenotypes of the two E3 ligases involved in Notch signalling, Neur and Mib1. While the loss of function of *mibl* causes the formation of well-separated supernumerary SOPs/bristles, the loss of *neur* causes the formation of small clusters of SOPs (weak neurogenic phenotype) [15–18]. Finally, the TFB model cannot explain why *neur* loss of function causes a weaker neurogenic phenotype than that of the typical neurogenic phenotype of the Notch pathway mutants [15]. Overall, these observations raise significant questions on the mechanism underlying the SOP selection and call for alternative models for this process.

Theoretical analyses of the TFB model show that CI could play an important role during lateral inhibition. In particular, CI could enhance the fidelity and speed of the lateral inhibition selection process by generating a sharp switch between signal-receiving and signal-sending states (reviewed in [19]). This is since an imbalance between Notch receptors and ligands at the cell surface can lead to a strong suppression of either the receptors or ligands, depending on the ratio between them. However, while CI has been identified as a regulatory mode during several developmental processes, experimental evidence for its requirement for SOP selection is missing.

We recently re-examined the formation of the SOP of the large mechanosensory bristle, the macrochaeta [15]. We found that three main factors regulate SOP selection: (i) The expression of proneural genes is broad and non-uniform across the notum with peak expression profiles around the future PNCs. (ii) There is a basal Notch activity within the entire notum driven by a combination of Mib1 activity and ubiquitin-independent activity. (iii) The combination between basal Notch activity and the peaks of the proneural gene expression profiles define a sub-group within the PNC from which SOP is selected in a neur-dependent manner (via lateral inhibition). The selected cell is typically at the peak position of proneural activity and initiates the expression of Neur. These observations suggest a picture where Notch has two distinct roles: to suppress differentiation across the notum and to generate lateral inhibition within a small sub-group of the PNC. Moreover, it suggests that SOP selection is mediated by regulation of Dl activity by Neur and Mib1 rather than transcription regulation of Dl (as suggested in the TFB model). Since in the TFB model the suggested role of CI during lateral inhibition is based on the feedback on Dl expression, it is unclear whether and how it affects SOP selection within this new picture.

Here, we use a combination of modelling and experiments to elucidate the role of CI during the selection process of the SOP. We develop a mathematical model termed the two channel SOP (TCS) model that explicitly incorporates regulation of Notch ligand activity by Mib1 and Neur, and accounts for CI between receptors and ligands. Using the unexpected observation that mammalian Dll1 has no cis-inhibitory activity in *Drosophila*, we experimentally test the prediction of the TCS model. We found that CI is essential for suppressing the basal Notch activity driven by Mib1. We identify a trade-off between the strength of CI and basal ligand activity mediated by Mib1, whereby reducing (strengthening) CI can be compensated by reducing (strengthening) basal ligand activity. Overall, our results provide evidence that CI is required for the selection of the SOP in the notum and the wing margin of *Drosophila melanogaster* and provide a new model for the SOP selection.

## Results

### Development of a mathematical model of SOP selection

To get a better understanding of the SOP selection in *Drosophila*, we developed a mathematical model, termed the two-channel SOP (TCS) model, that considers known regulatory interactions during the selection process (Figure 1A). This selection process is characterized by having two parallel channels of Notch signalling: one regulated and one unregulated. The regulated channel is based on ubiquitylation of Dl by Neur. We note that Neur can activate Dl even without ligand ubiquitylation [20], however, for simplicity we assume that activation is performed through ubiquitylation. We call this channel ‘regulated’ since Neur levels are indirectly inhibited by Notch signalling, forming a multicellular positive feedback characteristic of lateral inhibition [11]. The unregulated channel is based on Dl ubiquitylation by Mib1. We call this channel ‘unregulated’ since Dl ubiquitylation by Mib1 is not affected by Notch signalling. This unregulated channel leads to basal mutual inhibition normally preventing cells from differentiating into the SOP fate. Overall, the Dl ligand is then considered to be in one of three forms: Neur-, Mib1- or non-ubiquitylated. We note that although non-ubiquitylated Dl has been shown to have residual Mib1 independent activity [20], in the TCS model we simply account for Notch signalling only via ubiquitylated Dl and neglect Notch signalling from non-ubiquitylated Dl. We also assume that ubiquitylated ligands can get deubiquitylated at some finite constant rate, presumably by deubiquitylation enzymes. CI between Notch and Dl is also taken into account by assuming that Dl and Notch in the same cell can bind to each other forming an inactive complex, which is then removed from the cell surface [21]. The binding of Dl and Notch in the same cell reduces the levels of free ligands and receptors available for trans-activation, and therefore reduces Notch signalling. CI is assumed to occur for any of the ligand forms described above, namely for Neur-, Mib1- or non-ubiquitylated Dl [20].

**Fig. 1:**
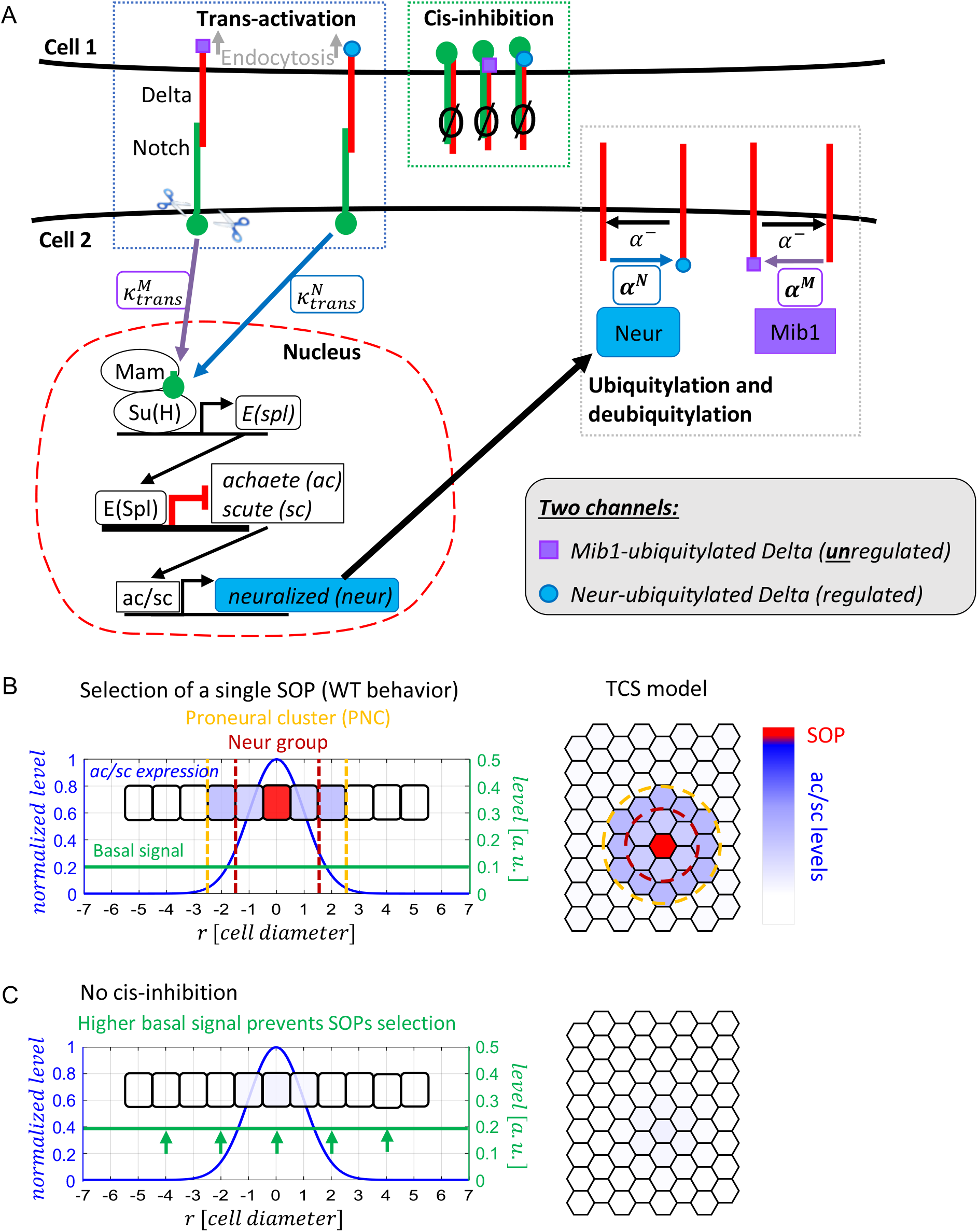
The TCS model predicts the role of CI during SOP selection. **(A)** Schematics of the TCS model. Blue frame: Trans-activation of Notch receptors in cell 2 by Mib1- or Neur-ubiquitylated Dl in cell 1, with respective trans-activation rates 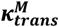 and 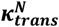. Red frame: transcriptional regulation of Ac/Sc and Neur by Notch activity. Gray frame: ubiquitylation of Dl by Mib1 (not regulated by Notch activity) and Neur (regulated by Notch activity), with respective ubiquitylation rates *α^M^* and *α^N^*. Mib1- or Neur-ubiquitylated Dl can get deubiquitylated at a rate *α*^-^. Green frame: CI between Dl and Notch in the same cell forms an inactive complex (0 sign), which prevents trans-activation via the cis-inhibited ligand and receptor. (B-C) Selection of SOP depends on the strength of CI. (B) Simulation result of the TCS model using the default parameters and a Gaussian profile for the Ac/Sc expression rate (blue line) and initial levels (See also methods, table S1 and Fig.1S1). A single SOP is selected at the top of the Gaussian (red cell). Cells in the Neur group are inhibited from becoming SOPs by Neur mediated signalling (lateral inhibition) from the selected SOP. Cells outside the Neur group, and inside the PNC, are inhibited from becoming SOPs by Mib1 mediated signalling (Basal signal, green line). Cells outside the PNC have a too low Ac/Sc expression to become SOPs (and are also inhibited by the basal signal). (C) Loss of CI, by setting the CI rate to zero (K_*c*_ = 0), leads to the loss of SOP selection due to an increase in basal Notch signal, mediated by Mib1. In this case the basal signal is increased by 80%, relative to the WT condition in (B).

For each protein described in the TCS model we assign a variable describing its level (concentration) in the cell, and for each dynamic reaction we assign a kinetic parameter, including the rates for ligand-receptor association/disassociation, trans-activation, genes expression, ubiquitylation, deubiquitylation, CI and degradation (See Methods and Tab. S1). We model the set of regulatory processes shown in Fig. 1A by a set of differential equations for the variables over time (as previously described in [21, 22]). These equations are then solved numerically and simulated on cells arranged in a fixed 2D hexagonal lattice (Figure 1B-C, Fig. S1).

Critical parameters of the system are the levels of the proneural proteins Ac and Sc (typically lumped together and denoted as Ac/Sc). The expression levels of Ac/Sc is known to be high in the proneural region defining the cells that belong to the PNC, while outside the PNC Ac/Sc levels are low [15, 23, 24]. Within the PNC, the central cell with the highest Ac/Sc levels is typically selected to become the SOP. The levels of Ac/Sc also link Notch signalling to Dl activity: Notch signalling downregulates Ac/Sc levels through the activation of E(spl), which in turn activates Neur (Fig. 1A) [17, 25]. In order to determine the parameter set and initial conditions to be used in our simulations, we initially analysed the system of equations in the area outside the PNC, where Ac/Sc expression rate is low, Neur is not expressed and SOPs cannot develop. We assume a uniform spatial steady state among all cells in this area, where only the unregulated channel of mutual Notch inhibition is functional. The parameters’ values gained from the uniform steady state analysis outside the PNC are then used in regions where ac/sc levels are high and SOPs can develop (Fig. S1A). In a condition where ac/sc levels are uniformly high, the TCS model results in a salt-and-pepper-like pattern of SOPs (Red cells in Fig. S1A’), typical for Notch mediated lateral inhibition process [11]. We note, though, that unlike standard models of lateral inhibition, the feedback here is not through regulating Dl expression levels (which remain constant) but rather by regulating Dl activity (i.e. Notch signalling induces expression of E(spl) that suppresses Neur, which in turn controls ubiquitylation of Dl). Thus, the model is consistent with the experimental observation that selection of SOP is normal, even when Dl is expressed under a constitutively active promoter [12].

We next wanted to simulate the situation where a PNC is defined. Experimental observations show that Ac/Sc activity is elevated within the future PNC (i.e. there is a pre-pattern of Ac/Sc) and decay away from the centre of the PNC. We therefore assume that Ac/Sc expression rate exhibits a decaying radial profile (Gaussian) from the centre of the PNC outwards (Fig. S1B-B’). A Gaussian profile of proneural activity was already previously suggested [15, 26]. The length scale of the decaying gradient is on the order of one cell diameter, consistent with observed PNC length scales. The profile of the Ac/Sc gradient is chosen such that a subgroup within the PNC, called the Neur group, accumulates high enough levels of Ac/Sc in order to participate in the selection of SOP via lateral inhibition. Naturally, the gradient also creates a bias such that a single central cell is selected as SOP from the Neur group, as it has the highest level of Ac/Sc and initiates the expression of Neur before its neighbours can do so [15, 26]. Outside the Neur group, only the unregulated Notch signalling channel is functioning. By tuning the parameters for the Ac/Sc expression rate inside the PNC, given the Gaussian profile of Ac/Sc, the TCS model results in a situation where only one SOP is selected from the Neur group (Fig. 1B). Thus, the TCS model is able to capture the wildtype (WT) case where a single SOP is selected from the PNC. In the coming sections, we show that the TCS model captures the previously observed mutant behaviours, as well as generate new predictions that are tested experimentally.

### The TCS model predicts that CI is required for SOP selection

We next tested if we can generate a prediction about the role of CI during SOP selection. To test the role of CI in SOP selection, we ran a simulation of the TCS model where CI rate is set to zero, but all other parameters remain fixed. This simulation shows a complete suppression of SOPs selection (Fig. 1C). The suppression occurs due to high basal Notch activity in all cells caused by the reduction in CI (see green line in Fig. 1B-C). Thus, the TCS model predicts that CI is required to set the basal Notch signalling to a level that defines the extent of the Neur group such that a single SOP is selected from the PNC.

### Identification of a ligand with no cis-inhibitory properties in *Drosophila*

In order to experimentally validate the prediction of the TCS model regarding the role of CI in SOP selection, we searched for a naturally existing ligand with no or weak cis-inhibitory abilities. Testing mammalian ligands, we found that murine Dll1 is a good candidate. To test the cis-inhibitory abilities of ligands one can express candidates in the wing primordium with *ptc*Gal4, which drives expression in a broad stripe of cells at the anterior side of the A/P compartment boundary of the wing imaginal disc as a gradient that increases to the posterior compartment boundary (Fig. 2A-A”). Dl-HA expressed in this gradient induces the ectopic expression of the Notch target gene *wg* in two parallel stripes in the dorsal and ventral compartment of the wing [27] (Fig.2B-B”). The common understanding of the formation of these two stripes is that expression of Dl leads to low Notch activity in regions of high expression due to CI, but high Notch activation in the adjacent posterior cells and also in anterior regions of low expression (Fig. 2B”). In agreement with the notion that CI depends on the ratio between Notch receptors and ligands, co-expression of Dl with Notch suppressed CI and leads to ectopic expression of Wg throughout the *ptc*Gal4 gradient (Fig. 2C).

**Fig. 2:**
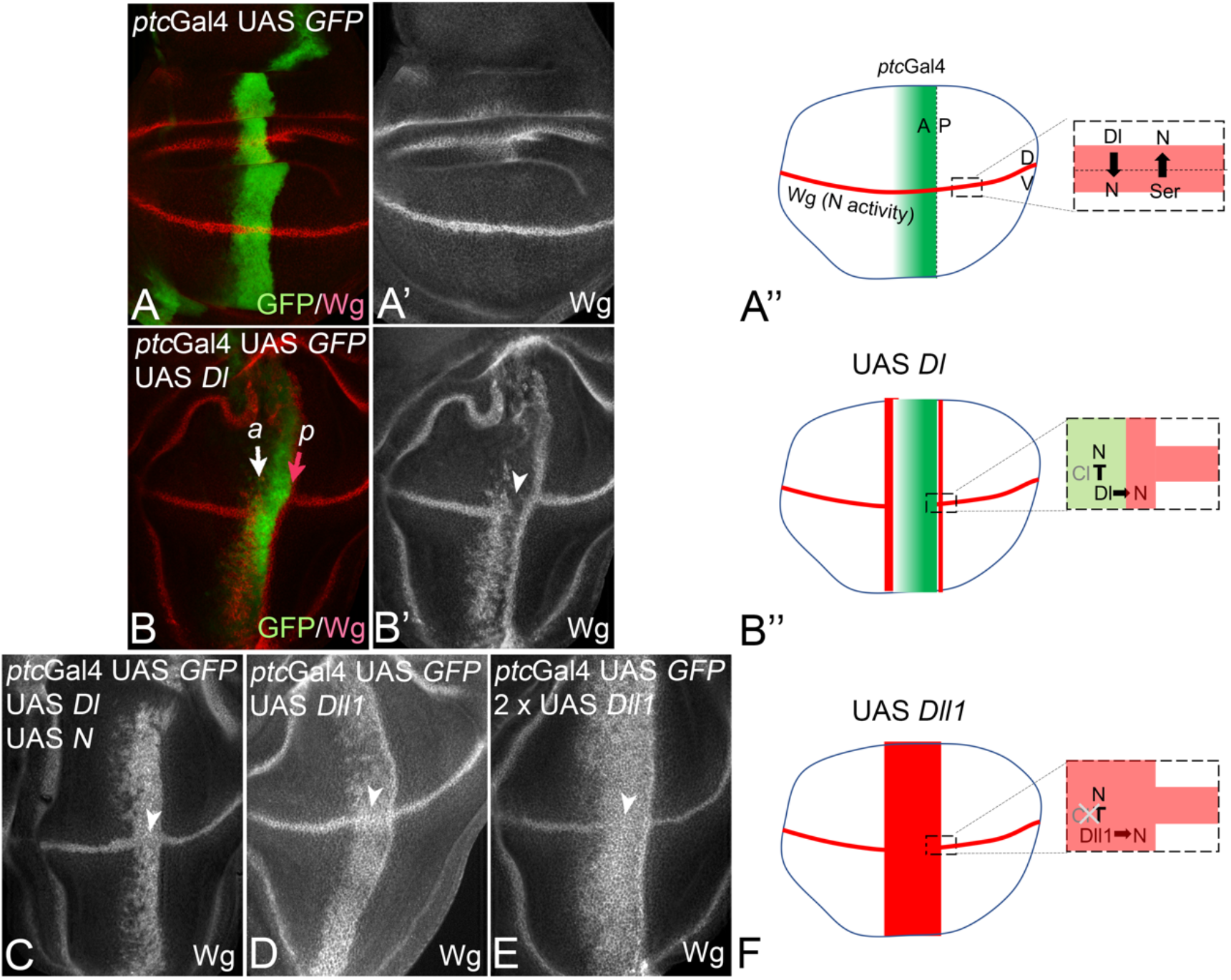
Dll1 has no cis-inhibitory abilities in Drosophila. **(A-A”)** *ptc*Gal4 drives expression of UAS constructs in a band along the anterior side of the A/P boundary (see A”). The expression is graded and decreases towards the anterior side. It intersects that of the stripe-like expression domain of Wg along the dorso-ventral (D/V) compartment boundary. **(B-B”)** Expression of Dl with *ptc*Gal4 results in the induction of two ectopic stripes of Wg expression perpendicular to its endogenous domain with no expression between them, which is a hallmark of CI. Of note, the ectopic expression of Dl suppresses signa**l**ling via CI also in the region where Wg is endogenously expressed (arrowhead). (C) Co-expression of Notch (N) with Dl suppresses CI and results in the ectopic expression of Wg throughout the whole *ptc*Gal4 domain. No suppression of the endogenous Wg domain is observed (arrowhead). (D-F) Ectopic expression of Dll1 in the *ptc*Gal4 domain does not lead to CI. In contrast to expression of Dl, the expression of one (D) or two (E) copies of Dll1 leads to a broad stripe of Wg expression, indicating the lack of CI (F).

We found that the expression of Dll1-HA also induced a broad band of ectopic Wg expression, similar to co-expression of Dl with Notch (Fig. 2D, F, compare with C). No region devoid of Wg expression was observed, neither in the ectopic, nor in the endogenous domains, indicating that Dll1 has little or no cis-inhibitory activity. We tested other Dll1 lines inserted at other genomic positions and found that they also lacked CI (not shown).

To test whether CI by Dll1 is eliminated, or only reduced, we co-expressed two Dll1 insertions to produce even higher levels of Dll1 expression (thus increasing the ratio between the Notch ligand and receptor [3, 21]). We found no evidence for CI even at these high expression levels of Dll1 (Fig. 2E).

Altogether, these results show that Dll1 possesses no or only very weak cis-inhibitory abilities in *Drosophila*. Therefore, we used Dll1 for the majority of our experiments. The expression of Dl-HA and Dll1-HA in MARCM clones in the wing primordium of the wing imaginal disc confirmed the same differences in the cis-inhibitory abilities of the ligands (Fig. 2S1D-G). While expression of Dl induces strong expression of Wg in cells adjacent to the clone, but much weaker expression within the clone (Fig. 2S1D, E), Dll1 induced strong levels of Wg expression throughout the clone and in adjacent cells (Fig. 2S1F, G). Note, that NICD expressing MARCM clones behave in a similar manner (Fig. 2S1H, I).

### CI is required for SOP selection in the notum and wing

To test whether the predictions of the TCS model regarding the requirement for CI for selection of the SOP (Fig. 1C), we employed the established assay where the Dl variants are expressed uniformly in *Dl Ser* mutant cell clones with the MARCM system [12]. As captured by the TCS model (Fig. 3B), the concomitant loss of *Dl* and *Ser* in PNCs of the notum, results in a strong neurogenic phenotype where many cells within the clone are selected for SOP development, indicated by the expression of the SOP marker Hindsight (Hnt) and the formation of tufts of bristles in the corresponding imago (Fig. 3A’-B’”). Ubiquitous expression of UAS *Dl* in the *Dl/Ser* mutant cells by *tubGal4* (MARCM clones) re-establishes the selection process and the bristle pattern in the imago (adult fly) [12, 20] (Fig. 3C-C’”).

**Fig. 3:**
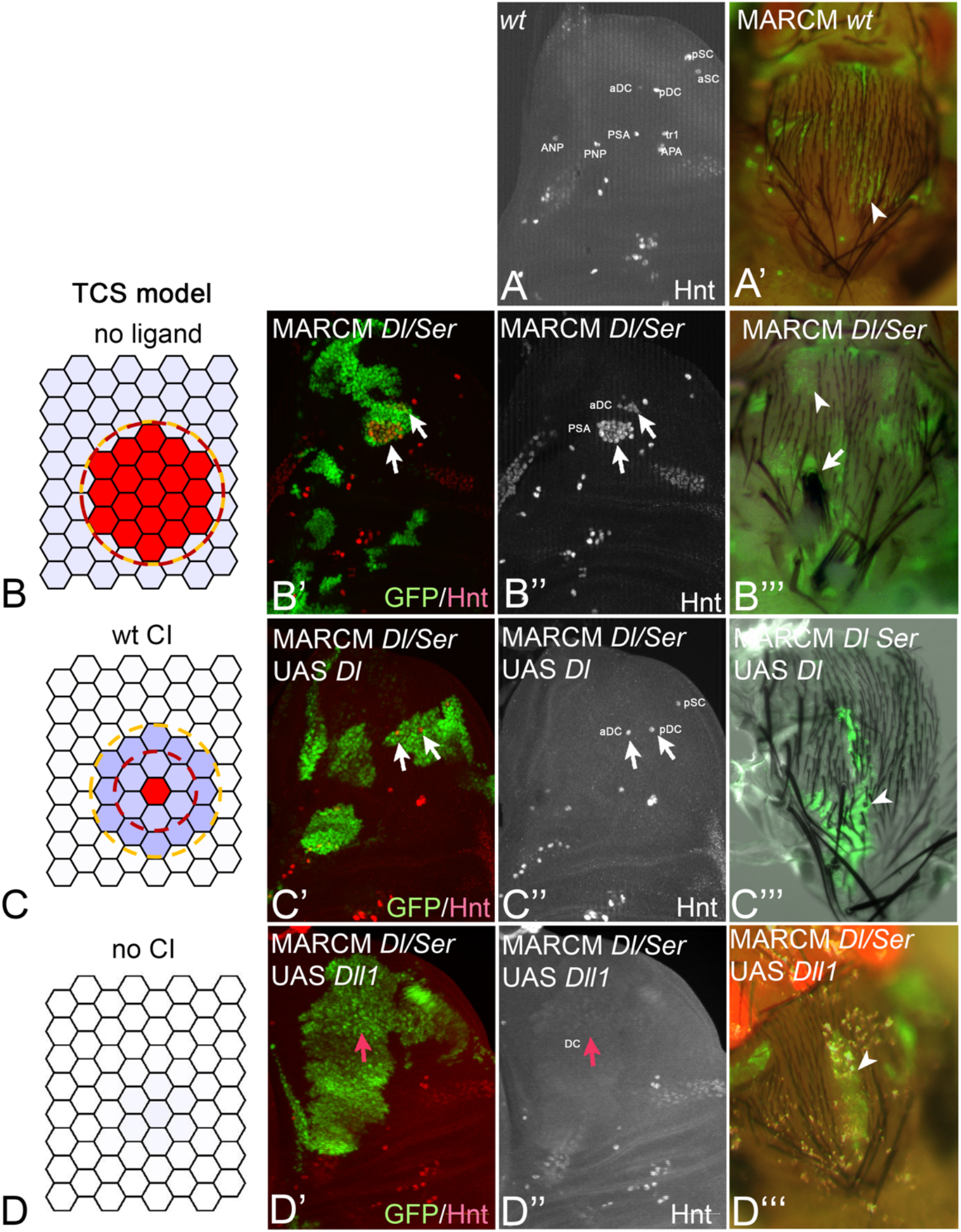
CI is required for the selection of the SOP in the notum. **(A-A’)** SOP pattern in a late third instar wing disc revealed by Hnt expression (A) and bristles of the adult wildtype notum (A’). GFP expression marks a MARCM clone (arrowhead). **(B-B’”)** The effect of *Dl/Ser* MARCM clones on SOP pattern. The TCS model (B) predicts a strong neurogenic phenotype in the absence ligands. Experiments with *Dl/Ser* MARCM clones (arrows in B’) show strong neurogenic phenotype within the clones in the wing disk (arrows in B”) and the formation of tufts of bristles in the case of the large sensory bristles (macrochaeta, arrow in B’”) and loss of the external parts of small sensory bristles (microchaete, arrowhead in B’”). **(C-C’”)** The effect of *Dl/Ser* MARCM clones expressing UAS-Dl on SOP pattern. The TCS model (C) predicts a single SOP when WT ligand (with CI) is uniformly expressed (same as Fig. 1B). Experiments with *Dl/Ser* UAS-Dl MARCM clones (arrows in C’) show normal SOP phenotype within the clones in the wing disk (arrows in C”) and in adult notum (arrowheads in C’”). **(D-D’”)** The effect of Dl/Ser MARCM clones expressing UAS-Dll1 on SOP pattern. The TCS model (D) predicts no SOP when a ligand with no CI is used (same as Fig. 1C). Experiments with *Dl/Ser*, UAS-Dll1 MARCM clones (red arrow in D’) show no SOP forming within the clones in the wing disk (red arrow in D”) and no bristles in the adult notum (arrowhead in D’”).

We next monitored the consequences of expression of the non-cis-inhibitory Dll1 for SOP development in two different developmental contexts: in the notum and in the wing. If not mentioned otherwise, the MARCM clone cells are double mutant for *Ser* and *Dl* to prevent any interference of the endogenous ligands.

#### Analysis in the notum

In contrast to UAS-*Dl*, the expression of UAS-*Dll1* in MARCM clones completely suppressed the formation of SOPs (Fig. 3D-D’”), confirming the prediction of the TCS model (Fig.1C). No Hnt-positive cells emerged in the clonal areas. The clones of freshly hatched or late pupal flies were devoid of bristles confirming that bristles development is abolished in the clone (Fig. 3D’”). The observed phenotype of Dll1 MARCM clones resembled that of expression of the activated form of Notch (NICD) in MARCM clones (Fig. 3S1A, A’).

To make sure that Dll1 induces Notch at least as much as Dl, we looked at the response of the Notch activity reporter Gbe+Su(H) in Dll1 and Dl MARCM clones known to have high sensitivity to Notch signalling (Fig. 3S1B-E’) [28]. We observed a strong increase in the expression of the Notch activity reporter in Dll1 MARCM clones in comparison to Dl-HA clones, indicating that Dll1 expression induces higher levels of Notch activity than Dl expression (compare Fig. 3S1C-C’ to Fig. 3S1D-D’). This conclusion was corroborated by co-expressing a *Notch-RNAi* construct along with Dll1 in *Dl/Ser* MARCM clones. In this case, we observed a strong neurogenic phenotype, similar to that observed for *Dl/Ser* MARCM clones, indicating that activation by Dll1 requires Notch receptors (Fig. 3S1E, E’).

Consistent with these experimental results, a simulation of the TCS model where both CI and Notch expression rates are set to zero results in the strong neurogenic phenotype (Fig. 3S1F). These results indicate that Dll1 activates Notch and that this activation leads to the suppression of SOP formation. Finally, to confirm that CI is required for the suppression of the basal Notch activity, we turned to a previously identified Ser variant that lacks CI [9]. In this variant, the sixth EGF repeat of its extracellular domain is deleted (SerΔEGF6). The expression of SerΔEGF6 in *Dl/Ser* double mutant MARCM clones also lead to suppression of SOP formation phenocopying the behaviour of Dll1 (Fig. 3S1G, G’).

In summary, the presented results support the prediction of the TCS model that ligands that lack CI cannot mediate SOP selection in the notum, indicating that CI is an essential requirement in this process. They also support the prediction that ligands that lack of cis-inhibitory activities induces elevated levels of the basal Notch activity that are not compatible with SOP development. A critical parameter for CI is the ratio between the ligand and Notch levels [3, 27]. We therefore aimed to suppress CI of the endogenous ligands during SOP selection by changing the ratio in favour of Notch. We expected that an increase in Notch expression level overcomes CI and elevates the basal Notch activity. To experimentally test this, we over-expressed UAS *Notch* in MARCM clones. This over-expression of Notch in wildtype cells led to an increase in activation of the Notch pathway, indicated by elevated expression of the Gbe+Su(H) Notch activity reporter (Fig. 3S2A, A’). It also suppressed SOP formation in the disc and bristle development (Fig. 3S2B-B”). The suppression by Notch elevation, required the presence of the endogenous ligands, since it was abolished in *Dl/Ser* mutant MARCM clones expressing Notch (Fig. 3S2C, C’). These results further confirm that the suppression of the cis-inhibitory abilities of the endogenous ligands results in a failure of SOP formation.

#### Analysis in the wing

The SOPs of some of the bristles of the anterior wing margin are determined during the late third instar stage in two stripes along the anterior half of the D/V boundary, adjacent to the domain of Wg expression (Fig. 4A-A”). Notch signalling has two roles in the determination of these SOPs: It induces the expression of Wg along the D/V boundary, which in turn induces the two stripe-like PNCs adjacent to it ([2, 29]; Fig. 4A”). In the established PNCs, Dl signalling is required for the selection of the SOP. Thus, the Notch pathway is indirectly (via induction of Wg expression) required to establish the PNC and subsequently to select the SOP within it. Ectopic SOP development can be induced in the wing blade by expression of Wg or Dl [29] (Fig. 4B-C”). In the case of ectopic Wg expression, the arising SOPs are regularly distributed and well-separated from each other (Fig. 4B-B”). Depletion of Notch function in Wg expressing MARCM clones resulted in a strong neurogenic phenotype in the clone, indicating that the regular spacing of the SOPs is achieved by Notch signalling (Fig. 4S1A-A”). Note, that this pattern of SOPs is predicted by the TCS model (Fig.1S1A’). MARCM clones that expressed Dl induced two PNCs, one stripe-like PNC outside the clone (outside PNC) and one less defined inside the clone (inside PNC). The PNCs give rise to well-separated SOPs that express Hnt and Neur (Fig. 4C-C”).

**Fig. 4:**
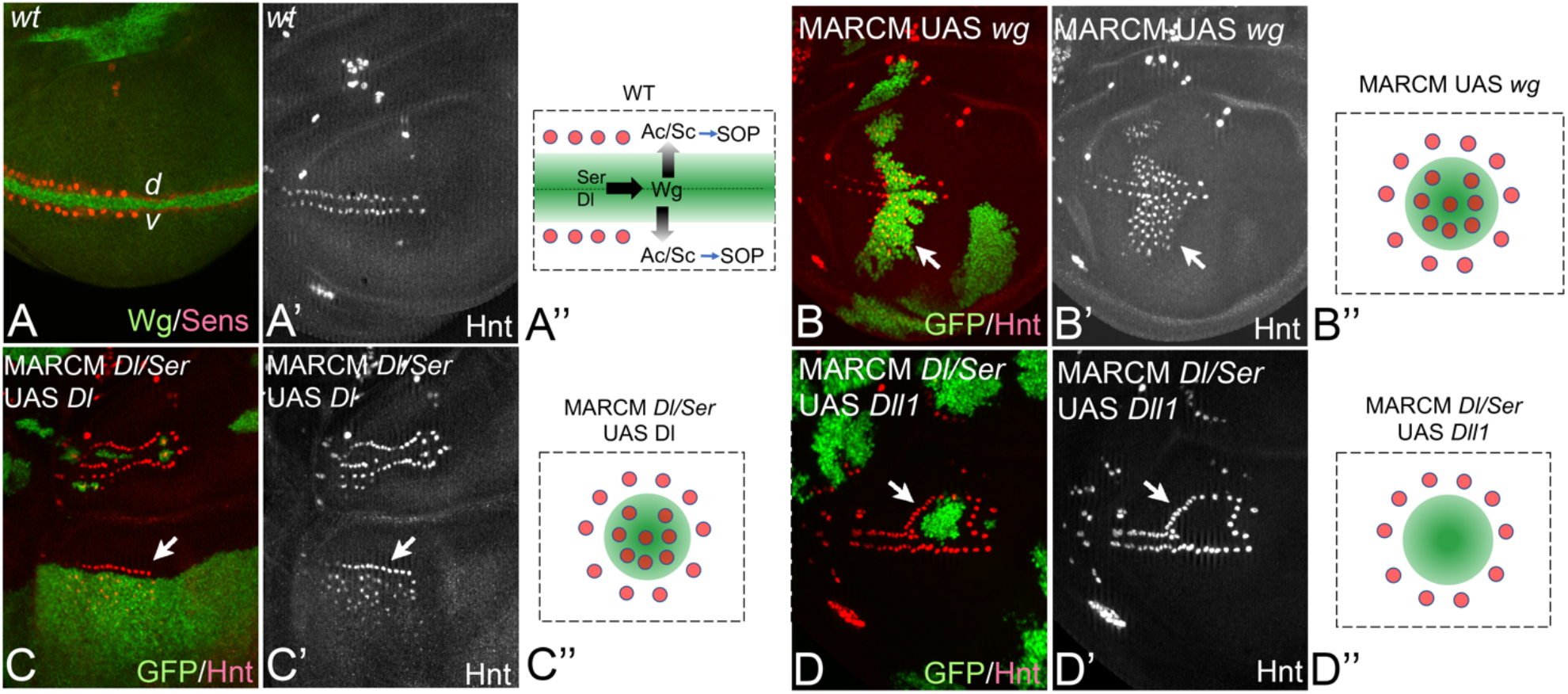
CI is required for the selection of the SOP in the wing. **(A-A”)** Induction of SOP formation along the D/V compartment boundary in the anterior compartment by Notch and Wg. The activity of the Notch pathway induced by mutual signalling of Ser and Dl establishes the expression of Wg along the D/V boundary. Wg secreted by the boundary cells induces the expression of Ac/Sc in adjacent dorsal and ventral cell to establish two stripe-like proneural cluster from which the SOPs are selected via Notch signalling. **(B-B”)** Ectopic expression of Wg via MARCM clones results in the establishment of ectopic proneural clusters and ectopic SOPs within the clone (inside PNC) and outside the clone (outside PNC, arrow). The SOPs are well-separated, because of the Notch mediated lateral inhibition process (see also Fig. 4S1A-A”). **(C-C”)** Expression of Dl in *Dl/Ser* MARCM clones induces SOP formation inside and outside the clone (arrow). **(D-D”)** In contrast to Dl, the expression of Dll1 in *Dl/Ser* MARCM clones suppresses SOP formation in the clone, but induces it outside the clone.

The expression of the activated form NICD resulted in the induction of high expression of Wg throughout the clone, but at the same time suppressed SOP formation in there (Fig. 2S1F, F’ and Fig. 4S1B-B”). However, regularly spaced SOPs were observed around the clone (arrow in Fig. 4S1B-B”). This finding indicates that continuous high Notch activity in the clone prevented SOP development. However, the induced Wg can establish the outside PNC, where SOP formation occurs normally. The same phenotype was observed upon expression of the non-cis-inhibitory ligand Dll1. It resulted in a failure of SOP formation inside the clone, but did not affect SOP development outside (Fig. 4D-D”). The suppression of SOP development inside the Dll1 expressing clone was relieved upon depletion of Notch (Fig. 4S1C-C”). A similar phenotype like Dll1, was induced by co-expression of Dl with Notch to suppress CI in MARCM clones (Fig.4S1D-D”). The findings confirm that CI is necessary to allow SOP formation to occur, also in the ectopically induced inside PNCs in the wing primordium.

### Ligands without cis-inhibitory abilities prevent the formation of the Neur-group

We next wanted to test whether CI is required for the formation of the Neur group. To do that, we first examined whether Neur expressing cells are observed in the presence or absence of CI in a MARCM clone setup. We observed Neur expressing cells in *Dl/Ser* mutant MARCM clones that express Dl, but not in clones that express Dll1, suggesting a Neur-group has formed in the Dl but not in the Dll1 expressing clones (Fig. 4S1E-F” and Fig. 5A-B’).

**Fig. 5:**
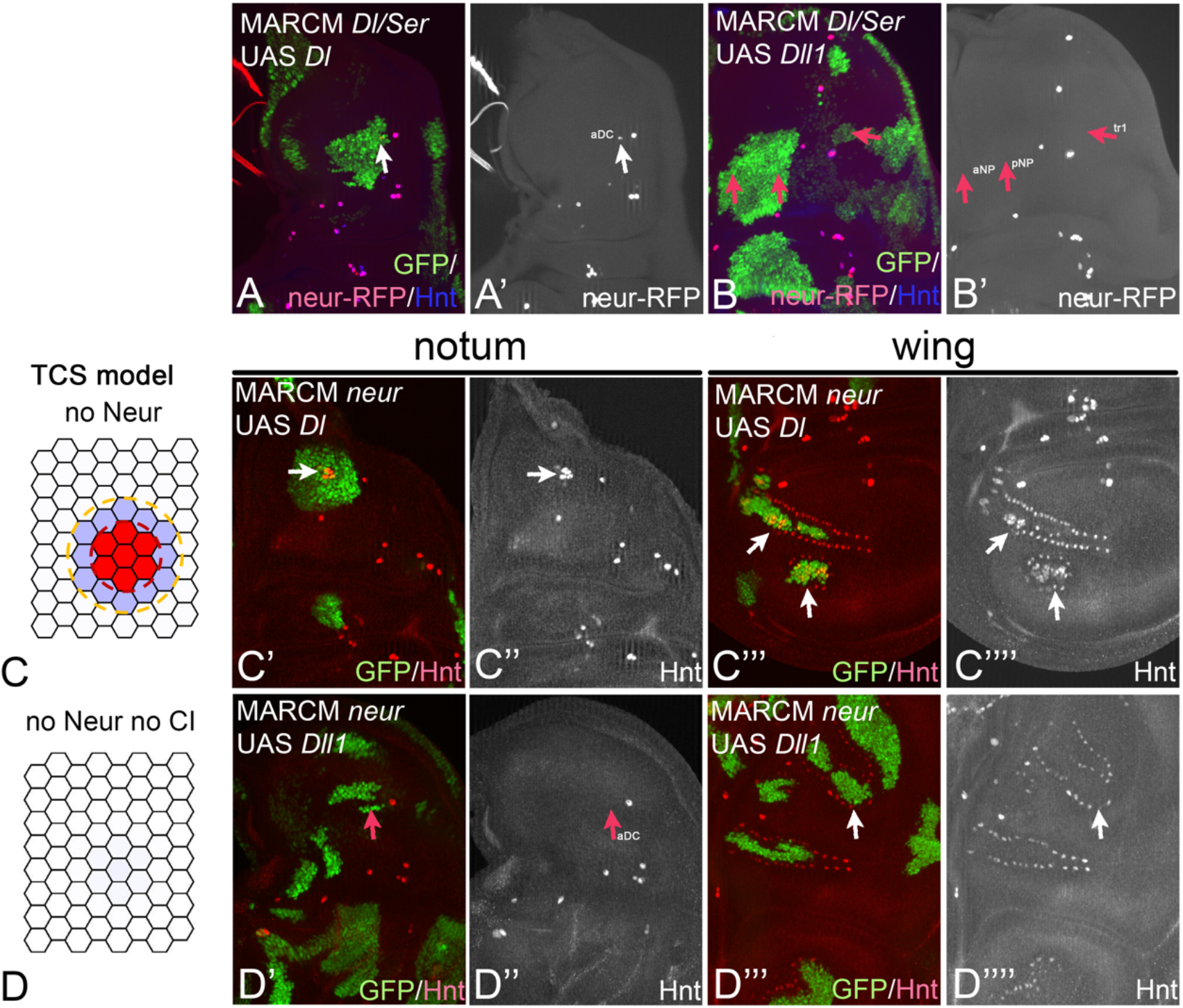
The formation of the Neur group requires CI. **(A-B’)** Expression of Neur in the future SOP is only observed in *Dl/Ser* MARCM clones that express a cis-inhibitory ligand. (A. A’) *Dl/Ser* MARCM clone expressing Dl shows a single Neur expressing SOP within the clone (arrow). (B, B’) Expression of Dll1 in *Dl/Ser* MARCM clones suppresses SOP formation in the clone (arrows) and Neur expression is activated in the clone cells. **(C-C”“)** The effect of *neur* MARCM clones (no Neur within the clone) expressing UAS *Dl* on SOP pattern. The TCM model (C) predicts that in the absence of Neur, a subgroup of cells within the PNC (the Neur group) will emerge when a WT ligand (with CI) is uniformly expressed. Experiments with Neur, UAS *Dl* MARCM clones in the notum (arrows in C’, C”) and in the wing disk (arrows in C’”, C””) show the formation of small groups of SOPs within the clones (weak neurogenic phenotype). **(D-D”“)** The effect of *neur* MARCM clones (No Neur within the clone) expressing UAS-Dll1 on SOP pattern. The TCM model (D) predicts that in the absence of Neur and with no CI, no SOPs will be formed. Experiments with Neur UAS *Dll1* MARCM clones in the notum (arrows in D’, D”) and in the wing disk (arrows in D’”, D””) show no SOPs within the clones. The result suggest that CI is required for the formation of the Neur-group.

It has been previously shown that Neur mutants lead to a weak neurogenic phenotype where a subset of PNC cells adopts the SOP fate [15, 18, 25]. This observation is captured in the TCS model with the same set of parameters used to simulate the WT case and the expression rate of Neur is set to zero (Fig. 5C). The subset of cells adopting SOP fate are expected to be those that express high enough levels of Ac/Sc to overcome the basal Notch activity provided by the non-regulated channel. Indeed, MARCM clones that lack *neur* function and express Dl showed the weak neurogenic phenotype in both the wing and the notum (Fig. 5C’-C””). In line with this observation is also that expression of a Dl variant where the Neur binding site is replaced by alanins (Dl-NxxN2A-HA, [20]) in *Dl Ser* MARCM clones, cannot mediate correct selection of SOPs. Instead, a weak neurogenic phenotype that resembles that of *neur* mutant clones was observed (Fig. 5S1A-B’). We next asked whether removing CI will suppress Neur expression in these cells. The TCS model predicted that in the absence of both Neur expression and CI (i.e. setting both the Neur expression and CI rates to zero in the model), the basal Notch activity would significantly increase, resulting in a decrease of Ac/Sc levels and in suppression of the weak neurogenic phenotype (Fig. 5D). This prediction was validated experimentally as *neur* mutant MARCM clones that express Dll1 showed no neurogenic phenotype in both the wing and the notum (Fig. 5D’-D””). This shows that, in contrast to ligands with cis-inhibitory abilities, CI-free ligands suppress SOP development in a Neur-independent manner. Combined with the finding that expression of Dll1 prevents the expression of Neur, these results support the notion that the basal Notch activity created by the CI free ligands is above the threshold that allows the definition of the Neur group and therefore allows the selection of a SOP fate.

### Enhanced CI results in an excess of SOPs

We next wanted to test what is the effect of enhanced CI on SOP selection. Analysis of the TCS model shows that increasing CI, without changing any other parameter, should result in a corresponding increase in the number and density of SOPs (Fig. 6A). This is because enhancing CI suppresses the basal Notch activity, which expands the domain of the Neur group within the PNC. Also, from the model, for strong enough CI (relative to the trans-activation channels) cells in the PNC become more refractory to trans-activation, which result in a denser SOPs pattern with directly neighbouring SOPs (See Fig. 6A “4 times wt CI”). In the wing primordium, it has been shown that Ser is more cis-inhibitory than Dl [3, 8]. We therefore monitored the effect of high CI on the selection of the SOP in the wing primordium. In agreement with the TCS model predictions, the ectopic expression of Ser in MARCM clones induced an excess of SOPs inside the ectopic PNCs (Fig. 6B-B”, arrow), indicating that Ser is unable to provide sufficient signalling activity for SOP selection within the clone. However, like Dl, Ser also induced the formation of well-separated SOPs outside the clone (Fig. 6B’, arrowhead). In contrast, expression of the CI-lacking SerΔEGF6 suppressed SOP formation inside, but not outside the clone (Fig. 6C-C”, arrowhead), suggesting that the high CI of Ser prevents the proper selection of SOPs in Ser expressing *Dl Ser* mutant MARCM clones. These results show that while reducing CI leads to loss of SOP, increasing CI drive the formation of extra, often more dense, SOPs.

**Fig. 6:**
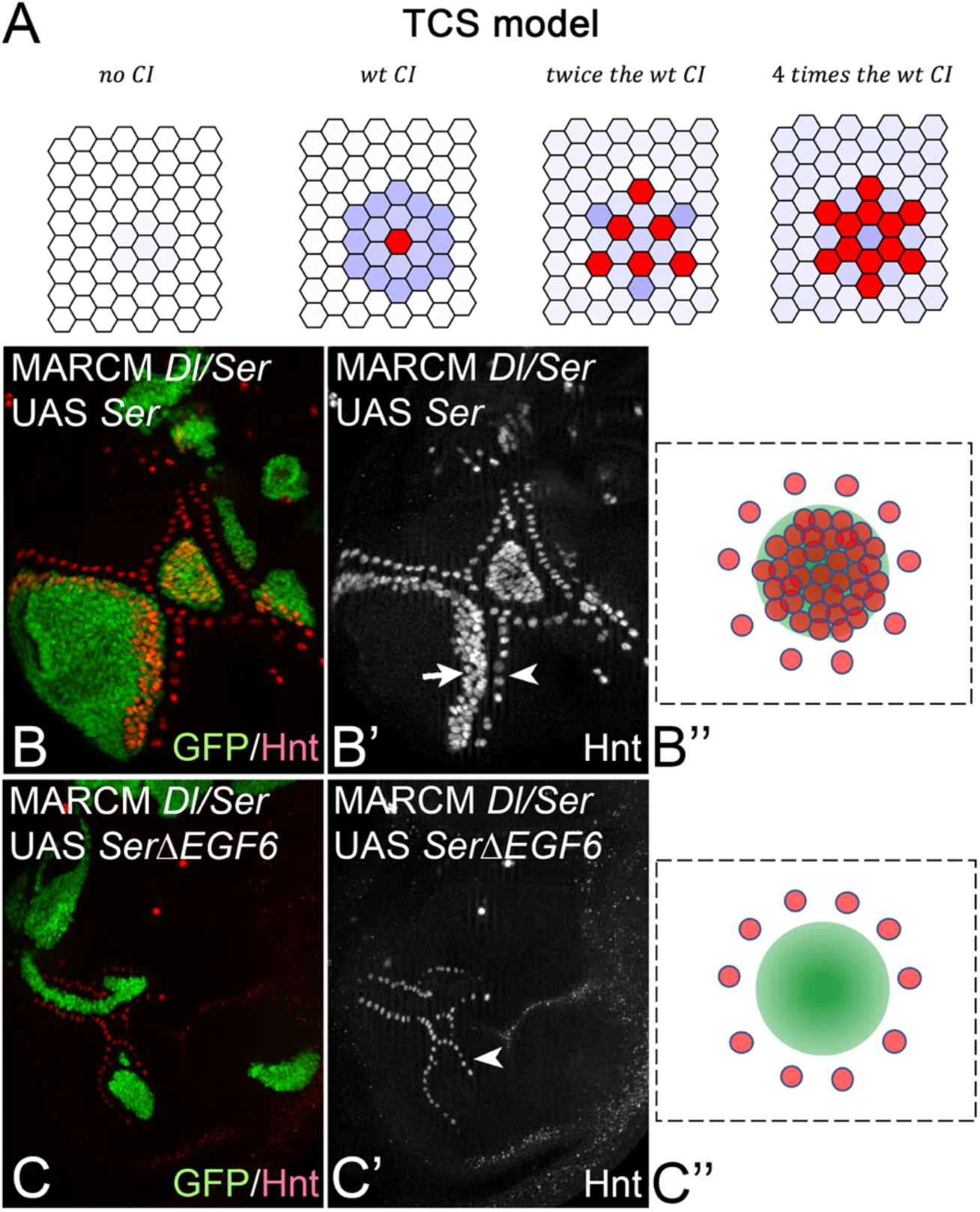
Ligands with strong CI promote formation of extra SOP. **(A)** The TCS model shows that increase in the strength of CI leads to the formation extra SOPs. The relative level of CI strength (defined by the parameter *K_c_*, see methods) compared to WT parameter are as indicated. **(B-B”)** Expression of Ser in *Dl/Ser* MARCM clones in the wing disc induces extra SOPs inside the clone (neurogenic phenotype, arrow) and normal SOPs outside the clone (arrowhead). **(C-C”)** In contrast, the expression SerΔEGF6, a Ser variant that lacks CI, suppresses SOP formation inside the clone completely. The SOPs outside the clone are still generated (arrowhead), indicating that SerΔEGF6 can signal in trans.

### A compensation mechanism between CI and the unregulated Notch channel

The overall picture that emerges from the modelling and experimental results is that a key aspect in SOP selection is the interplay between basal Notch signalling (mediated by the unregulated Mib1 channel) and CI which suppresses the basal Notch signalling to a level that allows the formation of the Neur group. To test the role of unregulated Notch signal, we have simulated in the TCS model the cases where Mib1 is over-expressed (higher unregulated channel strength, Fig. 7A) and where Mib1 is removed (lower unregulated channel strength, Fig. 7B). The TCS model predicted that higher level of Mib1 should lead to suppression of SOP due to higher basal Notch signalling (Fig. 7A), while lower levels of Mib1 should lead to formation of extra SOP due to lower basal Notch signalling (Fig. 7B). These predictions match the experimental observations as over-expression of Mib1 in MARCM clones lead to suppression of some of the SOP (Fig. 7A’-A”), while a knockout of *mib1* leads to formation of ectopic well separated SOPs (Fig. 7B’-B”). We note that we have previously shown that Notch signalling still persists (albeit, at a lower level) even if *mib1* is knocked out [20]. We therefore assumed in the simulation that the unregulated Notch channel is reduced but does not vanish when we knock out *mib1* (see methods).

**Fig. 7:**
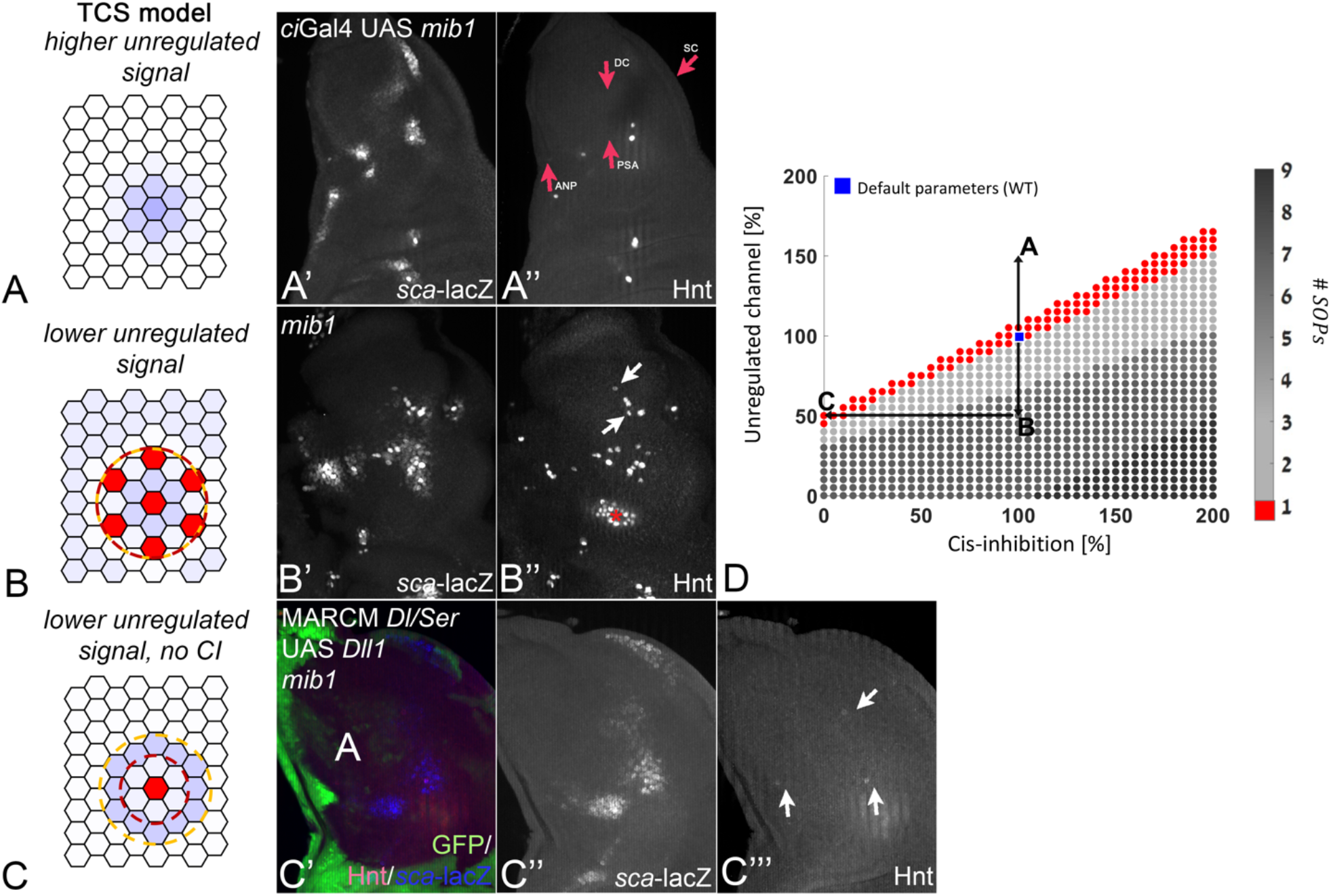
A trade-off between CI and the unregulated Notch channel. **(A-A”)** The effect of Mib1 over-expression on SOP phenotype. The TCS model (A) predicts that an increase of 50% in the activity of the unregulated channel, corresponding to higher Mib1 expression, should lead to suppression of SOP. Experiments where Mib1 was over-expressed showed that while PNCs are still defined (marked by *sca*-LacZ in A’), some of the SOPs within these PNCs are suppressed as predicted by the model (arrows in A”). **(B-B”)** The effect of *mib1* knockout on SOP phenotype. The TCS model (B) predicts that a reduction of 50% in the activity of the unregulated channel, corresponding to a reduced (but not completely suppressed) activity due to Mib1 knockout, should lead to the selection of more than one SOP from the PNC. Experiments where *mib1* function was deleted showed that PNCs are still defined (marked by *sca*-LacZ in B’), yet a few extra SOPs emerge within the clones, as predicted by the model (arrows in A”). **(C-C’”)** The combination of *mib1* knockout and removal of CI leads to normal SOP selection. The TCS model (C) predicts that a reduction of 50% in the activity of the unregulated channel (as in B) together with suppression of CI, should regain the WT phenotype of a single SOP. This prediction was realized experimentally by looking at *Dl/Ser* UAS *Dll1* MARCM clones (the region that lacks GFP in C’) in the background of a *mib1* knockout. As predicted experimentally, PNC clusters within the clone (marked by *sca*-LacZ in C”) showed the emergence of a single SOP within each clones (arrows in C’”). **(D)** A phase diagram showing the TCS model predictions for a range of parameters. The x-axis values refer to the relative levels of CI strength, *K_c_*, with respect to the value used to simulate the WT. The y-axis values refer to the relative levels of the ratio between unregulated and regulated Notch activity, 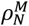, with respect to the value used to simulate the WT (see methods). The number of SOPs for each parameter set are shown by the colour bar. The value used for simulating the WT (Blue square) and the mutant cases shown in (A), (B), and (C) are indicated.

Next, we wanted to test the interplay between CI and the unregulated Notch channel. The TCS model predicted that the formation of ectopic SOPs in *mib1* knockout can be rescued by the loss of CI in the non-cis-inhibitory ligand, which promotes higher basal Notch signalling (Figure 7C). This situation was simulated by setting both the CI rate to zero and reducing the ratio between Mib1 and Neur ubiquitylation by half. The TCS model in this case predicts the formation of a single SOP. We experimentally tested this prediction from the model by looking at *Dl Ser* MARCM clones in a *mib1* mutant background that express a non-cis-inhibitory Dll1. As predicted by the TCS model, single SOPs appear within the domains of the clones, confirming the compensation mechanism between *mib1* activity and CI (Fig. 7C’-C’”).

To better understand the interplay between the unregulated Notch channel and CI, we have performed a comprehensive phase space analysis of the TCS model on the dependence of the SOP phenotype on the two key parameters of the system: The strength of CI and the relative strength of the signal from the unregulated channel (Fig. 7D). In this analysis, the strength of CI is described by the level of CI rate compared to the WT value. The parameter of the relative strength of the unregulated channel is described by the ratio between ligand activity associated with ubiquitylation by Mib1 and the maximal ligand activity associated with ubiquitylation by Neur (see methods). The phase space analysis provides a comprehensive view of the trade-off between CI and the unregulated Notch channel. Decreasing or increasing the strength of CI (moving horizontally on the Fig. 7D) would lead to a decrease or increase in the number of SOP, respectively (corresponding to Fig. 3C-D’” and Fig. 6). Similarly, increasing or decreasing the strength of the unregulated Notch channel (moving vertically on the Fig. 7D, points A and B) would lead to a decrease or increase in the number of SOP, respectively (corresponding to Fig. 7A-B”). Finally, decreasing both CI and the unregulated Notch channel, would lead to the WT SOP phenotype (point C in Fig. 7D, experimentally observed in Fig. 7C’-C’”). Thus, the phase space analysis highlights the conclusion that a selection of a single SOP relies on the trade-off between Notch signalling from the unregulated channel and CI.

## Discussion

The selection of the SOP is a classical, extensively studied, process mediated by Notch signalling where a single cell is selected from an equivalence group, the PNC. This process has been considered as a prime example of selection by lateral inhibition. Here, we propose a new model, the TCS model, for the selection of a single SOP from the PNC, which significantly differs from the classical models of lateral inhibition (referred to as the TFB models). The new aspects of the TCS model include: (i) The Notch ligand activity, rather than the ligand expression level, is regulated by Notch signalling. (ii) There are two distinct channels that control Dl activity associated with the E3 ubiquitin ligases Neur and Mib1. While the *neur* channel is regulated by Notch activity, the *mib1* is not regulated by Notch activity and produces a broad basal Notch activity across the notum. (iii) The PNC is defined by gradients of expression of the proneural genes *ac* and *sc*. Within the PNC, the subgroup that participates in the *neur*-controlled lateral inhibition process (the Neur group) is defined by both the gradient profile and the basal Notch activity driven by *mib1*. (iv) CI is required for the suppression of basal Notch activity and determines the extent of the Neur group within the PNC.

The unexpected discovery that mammalian Dll1 does not cis-inhibit Notch in *Drosophila* allowed us to show for the first time that CI is required for SOP selection and to test critical predictions of the TCS model. The TCS model can account for two earlier observations that could not be explained by the TFB model: (i) That unregulated homogenous expression of Dl can also mediate SOP selection [12, 13]. And (ii) that loss of *ac* and *sc* did not affect Dl expression [14, 15].

The TCS model also accounts for the distinct mutant phenotypes of *Dl/Ser* (neurogenic phenotype)*, neur* (weak neurogenic phenotype), and *mib1*. While in the Dl/Ser mutant all the cells in the PNC become SOP since both channels *(neur* and *mib1)* are removed, in the *neur* mutant only the cells in the *neur* group become SOPs since basal Notch activity through *mib1* remains and define the boundaries of the *neur* group. The *mib1* mutant leads to extension of the *neur*-group and consequently leads to the formation of multiple, well separated, SOPs. This is since *neur*-mediated Notch activity still exists and lateral inhibition is not restricted any more to a small sub-group.

Thus, the TCS model can explain the previously unexplained observations and captures the different Notch pathway phenotypes. It does so, with a single set of parameters highlighting the robust nature of the proposed model. We note that, our model relies only on Notch signalling between direct neighbours and does not invoke long range signalling by filopodia. The selection of only one SOP is achieved by restricting the potential to be selected for an SOP to a smaller sub-group of the PNC through Mib1-mediated basal Notch activity. This does not rule out a contribution of filopodia mediated Notch signalling in other process [30].

What is the role of CI in SOP selection? The overall picture that emerges from the TCS model and the experiments is that there is a trade-off between *mib1-mediated* basal Notch activity and the strength of CI. Without CI (replacing Dl by Dll1 or SerΔEGF6), there is a higher level of basal Notch activity, which completely abolishes the formation of SOP. With stronger CI, (replacing Dl by Ser) basal Notch activity is reduced and extra SOP emerge. Finally, by removing both CI (which increases basal activity) and *mib1* (which decreases basal activity) we get almost full compensation and regain a single SOP. Thus, CI is crucial for adjusting the level of basal Notch activity to determine the extent of the *neur* group and for the selection of a single SOP.

We would like to point out that CI could still directly contribute to the lateral inhibition process by making sender cells more refractory to signals from their neighbours (as has been suggested by [21]). The TCS model further suggests that for strong enough CI this refractory behaviour would lead to a denser SOP pattern (Fig. 6A). Consistent with this expectation, replacing *Dl* with the strong cis-inhibitory Ser ligand exhibit such denser SOP pattern (Fig. 6B).

The presence of multiple channels of signalling is not restricted to the SOP. For example, during differentiation of hair cells in the mammalian organ of Corti, signalling mediated by the Notch ligands Dll1 and Jag2 are involved in a lateral inhibition process, while signalling through Jag1, which acts in parallel, has different roles [31, 32]

## Acknowledgement

We thank Stefan Kölzer for excellent technical assistance. The Bloomington Drosophila Stock Center (NIH P40OD018537), the Vienna Drosophila Resource Center, the Drosophila Genomics Resource Center (NIH 2P40OD010949-10A1) and the Drosophila Hybridoma Bank supplied fly stocks and reagents.

## Funding

The work of TT and TK are supported by the Deutsche

Forschungsgemeinschaft (DFG) through Sachbeihilfe GZ: KL 1028/9-1

DS and TK are greatful for the the support of the DFG via Sachbeihilfe (Middle East Grant) KL1028/13-1

## Methods and Materials

### *Drosophila* methods

#### Fly strains

UAS *Ser-HA*, UAS *Dl-HA*, UAS *Dl-NEQN2A-HA* [20], UAS *Dll1-HA* [33], UAS *NICD* [13], UAS *SerΔEGF6* [9], UAS N-LV [34], UAS *N-RNAi* (Bloomington stock centre:BDSC_7078)), UAS *wg^E1^* [35], UAS *mib1* [36], *Dl^rev10^* e *Ser^RX82^* FRT82B [37], *neur^1^* e FRT82B [38], *mib1^EY09870^* [36], Gbe+Su(H)-lacZ [28], *sca*-lacZ (Bloomington stock centre, BSC5403), *neur*-RFP [39], *ptc*GAL4 [40], *ci*GAL4 [41]

##### Stocks for MARCM analysis

yw *hs*Flp *tub*GAL4 UAS GFPnls; ; FRT82B *tub*GAL80 /TM2,

yw *Ubx*Flp *tub*GAL4 UAS GFPnls; ; FRT82B *tub*GAL80 y+ / TM6 (Bloomington Stock Centre)

yw *Ubx*Flp; *ci*GAL4/CyOTb1; *mib1^EY09870^* FRT82B *ubiGFP tub*GAL80 / TM6B

#### Clonal Analysis

Clones were induced at the first larval instar stage (24–48 hr after egg laying) by heat shock (*hs*Flp, 1h at 37°C), or by *UbxFlp*.

#### Antibody Staining and Imaging

Antibody staining was performed according to standard protocols (Klein, 2006). Antibodies used: anti-Wg (4D4) (DSHB Iowa RRID:AB_528512), anti-ß-Gal (Cappel/MP Biomedicals RRID:AB_ 2313831) anti Hnt (1G9) (DSHB RRID:AB_528278), anti ELAV (7E8A10) *(DSHB* RRID:AB_528218), anti-futsch (22C10) (DSHB RRID:AB_528403). Alexa-Fluorochrome-conjugated secondary antibodies were purchased from Invitrogen/Molecular Probes. Images were acquired with a Zeiss AxioImager Z1 Microscope equipped with a Zeiss Apotome / Apotome2.

##### Model

For the full derivation of the TCS model please see supplemental material. This derivation includes a detailed description of the reactions in Figure 1A, formalization of these reactions in a set of differential equations, a steady state analysis and assumptions to extract the parameters of the model, and finally a description of the computer simulation and initial conditions. See also table S1 for the summary of the dimensionless variables and parameters that are used in the simulations.

The final set of dimensionless dynamic equations that are solved and simulated are: Ligands and receptor levels

###### Ligands and receptor levels

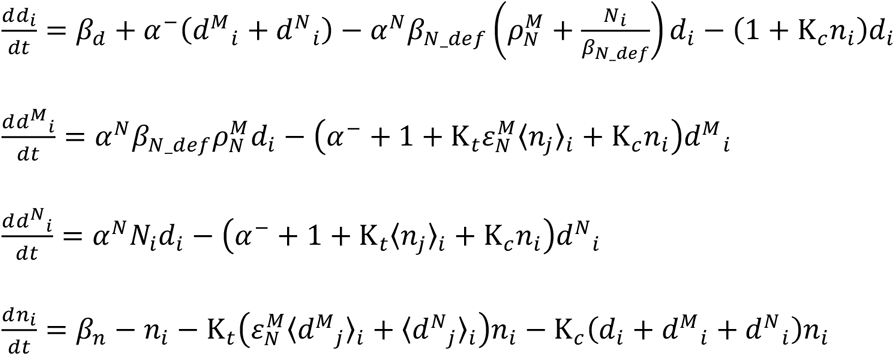

###### Intracellular components levels

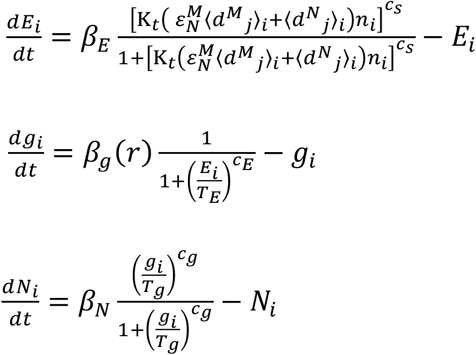

Where:

*d_i_, d^M^_i_, d^N^_i_, n_i_, E_i_, g_i_* and *N_i_* are the levels in cell *i* of non-ubiquitylated Dl, Mib1-ubiquitylated Dl, Neur-ubiquitylated Dl, Notch, E(Spl), ac/sc and Neur, respectively.

〈*d_j_*〉_*i*_, 〈*d^M^_j_*〉_*i*_, 〈*d^N^_j_*〉_*i*_, and 〈*n_j_*〉_*i*_ are the sum of the levels from cells *j* on the boundaries with cell *i* of non-ubiquitylated Dl, Mib1-ubiquitylated Dl, Neur-ubiquitylated Dl and Notch, respectively.

*r* is a spatial variable defining the gradient profile of ac/sc.

*β_d_*, *β_n_*, *β_E_*, *β_g_* and *β_N_* are the expression rates of Dl, Notch, E(Spl), ac/sc and Neur, respectively.

*β_N_def_* is the *β_N_* in the default set of parameters (wt condition).

*α^N^* and *α*^-^ are the rates of ubiquitylation of Dl by Neur and deubiquitylation of Mib1-or Neur-ubiquitylated Dl, respectively.

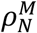 is the ratio between ligand activity associated with ubiquitylation by Mib1 and the maximal ligand activity associated with ubiquitylation by Neur.

K_*c*_ and K_*t*_ are the rates for CI and trans-activation, respectively.

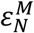 is the ratio between trans-activation of Notch from Mib1- and Neur-ubiquitylated Dl.

*c_s_*, *c_E_* and *c_g_* are hill coefficients associate with the levels of Notch activity ([*NICD,Mam,Su*(*H*)] complex), E(Spl) and ac/sc, respectively.

*T_E_* and *T_g_* are Hill half occupation levels associate with the levels of E(Spl) and ac/sc, respectively.

The equations are solved numerically, using a standard Matlab ordinary differential equations (ODE) solver, and the solution is simulated using a custom code given in: https://github.com/Udi-Binshtok/SOP_model_2021

## Supplemental Material

### Derivation of the TCS model

To quantitatively describe the selection of a single SOP from a PNC, during development of the wing imaginal disc of *Drosophila*, we developed a new model, termed the two channel SOP (TCS) model, that is based on a model previously described in [21]. The TCS model expands the classical model of lateral inhibition (reffered to as the TFB model, in the main text), first by taking into account two parallel channels of Notch signaling: a regulated channel (mediated by Neur) and an unregulated channel (mediated by Mib1) (Figure 1A); Second, by taking into account a spatial gradient for ligand activity (as described in more detail in the computer simulation part); and third, by accounting for CI between receptors and ligands. For simplicity, we only account for the Delta ligand (i.e. Neglected the contribution of Serrate).

#### Notation

The model describes the levels of Notch and Delta, as well as the levels of other components of the system including ligases Mib1 and Neur, ubiquitylated Delta (by Mib1 or by Neur), signal (activation complex) and its downstream genes (Figure 1A), in each cell *i* in a cell lattice. The following notation is used to describe the levels of these components:

*n_i_* ≡ *Notch levels in cell i*
*d_i_* ≡ *Delta levels in cell i*
*d^M^_i_* ≡ *Mib*1 *mediated ubiquitylated Delta levels in cell i*
*d^N^_i_* ≡ *Neur mediated ubiquitylated Delta levels in cell i*
*s_i_* ≡ [*NICD,Mam,Su*(*H*)] *complex levels in cell i*
*E_i_* ≡ *E*(*spl*) *levels in cell i*
*g_i_* ≡ *ac/sc levels in cell i*
*N_i_* ≡ Neur levels in cell i
*M_i_* ≡ *Mib*1 levels in cell i
〈*d_j_*〉_*i*_ ≡ *sum of Delta from cells j, on the boundaries with cell i*
〈*N_j_*〉_*i*_ ≡ *sum of Notch from cells j, on the boundaries with cell i*
[*n_i_d_j_*] ≡ *complex of Notch from cell i and Delta from neighbor cell j*

#### Mechanism

The interaction of the components within cell *i* and between cell *i* with neighboring cells *j* are described by Michaelis-Menten reactions [42]:

##### Delta ubiquitylation and deubiquitylation

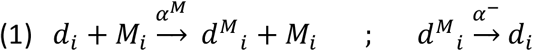

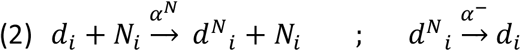

Where *α^M^* and *α^N^* are Mib1 or Neur mediated ubiquitylation rate, respectively, and *α*^-^ is deubiquitylating rate via an enzyme at a constant level.

##### Notch-Delta complexes and trans-activation

In the model we assume that any form of the ligand (non-, Mib1- or Neur-ubiquitylated ligand) can bind to the Notch receptor, but only the ubiquitylated forms can transduce a signal.

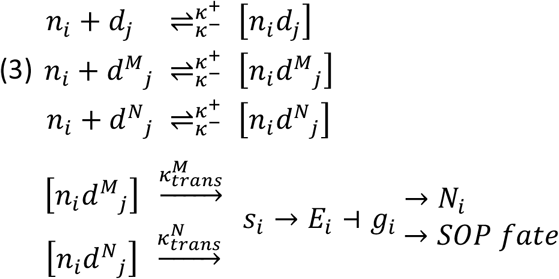

Where *κ*^+^and *κ*^-^ are the association and dissociation rates, respectively, between the free Notch and Delta state and the Notch-Delta complex state. 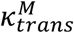 and 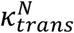 are the trans-activation rates (namely, the inverse time it takes for the NICD to get cleaved once it interacts with Delta) from Mib1- or Neur-ubiquitylated Delta, respectively.

##### Cis-inhibition

During CI Notch receptors and ligands in the same cell bind and create an inactive complex. The complex could dissociate back into the free receptors and ligands or go through degradation. In the model CI is considered for each of the ligand forms (non-, Mib1- or Neur-ubiquitylated Delta).

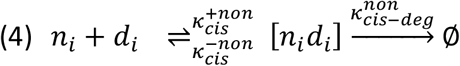

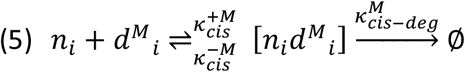

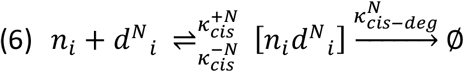

Where 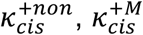 and 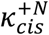 are association rates, 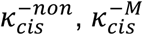 and 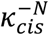 are disassociation rates and 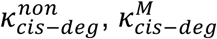 and 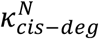 are degradation rates, of cis-inhibited Delta-Notch complexes from non-, Mib1- or Neur-ubiquitylated Delta, respectively. 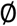 indicates degradation (no further reaction).

#### Equations

The activation and repression reactions in equation (3) (Figure 1A) are described in terms of Hill functions [43]. The reactions above are then converted into a set of ordinary differential equations (ODE) describing the change in the levels of the components in each cell *i* over time *t*:

##### Ligands and receptor levels

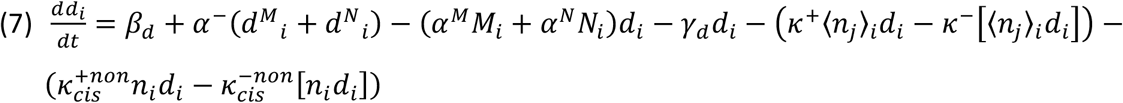

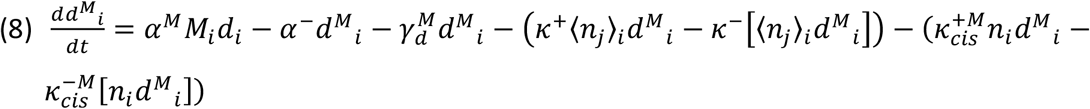

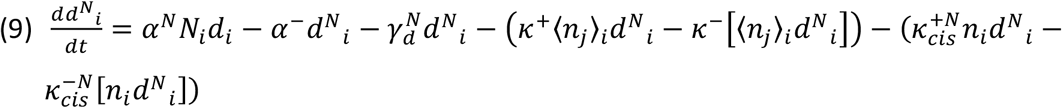

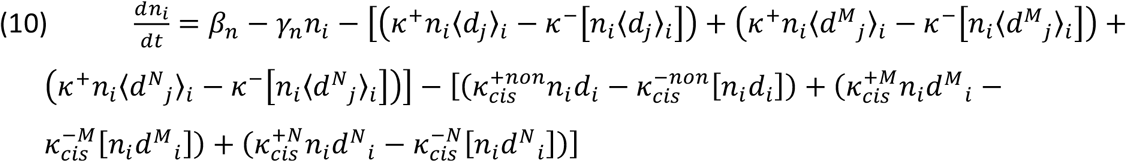

##### Delta-Notch complexes and signal levels

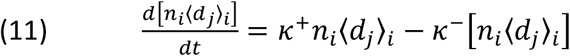

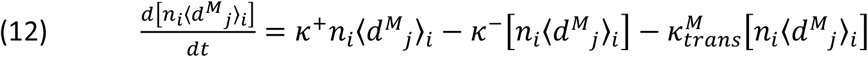

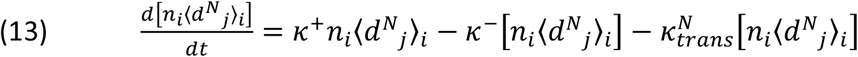

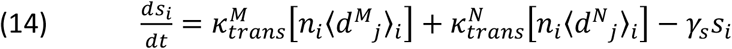

##### Intracellular components levels

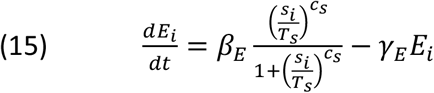

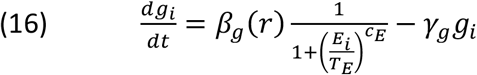

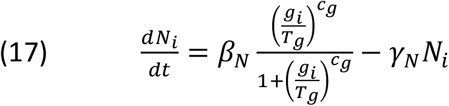

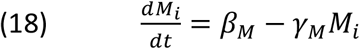

##### Cis-inhibition complexes levels

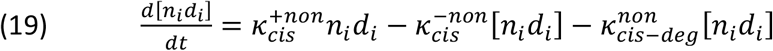

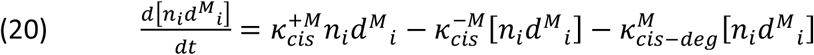

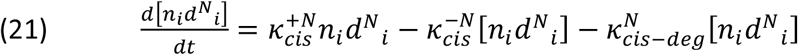

Where *β_x_* and *γ_x_* are the production and degradation rates of variable *x* (*x* = *n,d*,…), respectively. *r* is a position variable. *T_x_* and *c_x_* are the Hill function parameters: half occupation (effective dissociation Konstant, K_*d*_) and Hill coefficient, respectively, associated with the Hill function of variable *x*.

Note that we also term the half occupation *T_x_* as the *x threshold* level for promoting or inhibiting the expression of the variable that depends on *x*:

*T_s_* is the [*NICD,Mam,Su*(*H*)] (*s_i_*) threshold level for promoting *E*(*spl*) (*E_i_*) expression.
*T_E_* is the *E*(*spl*) (*E_i_*) threshold level for inhibiting *ac/sc* (*g_i_*) expression.
*T_g_* is the *ac/sc* (*g_i_*) threshold level for promoting *Neur* (*N_i_*) expression.

#### Simplification

In order to reduce the number of equations and parameters (equations (7) to (21)), we use the next set of assumptions and definitions.

1. We assume that cis-inhibited Delta-Notch complexes do not disassociate [21] and that the association rate is the same for all the Delta variations:

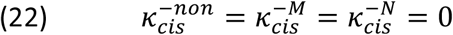

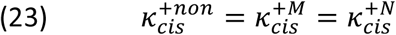
2. We assume a steady state approximation for Delta-Notch complexes, [*NICD,Mam,Su*(*H*)] complexes, and Mib1 levels:

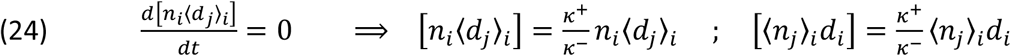

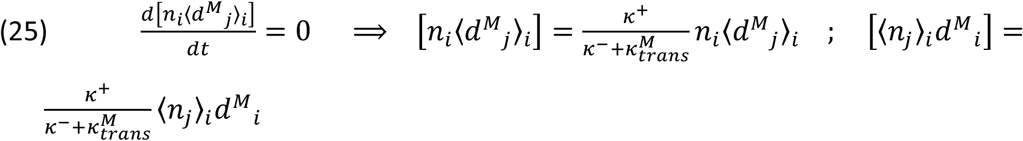

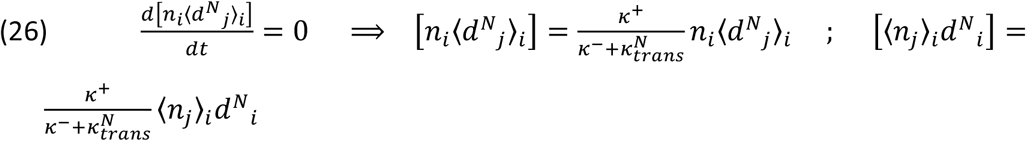

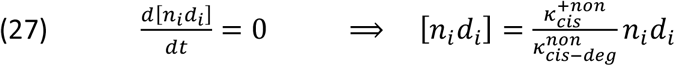

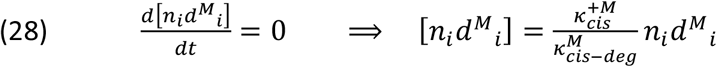

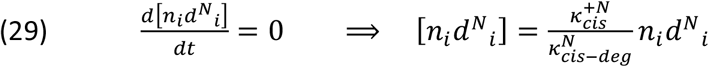

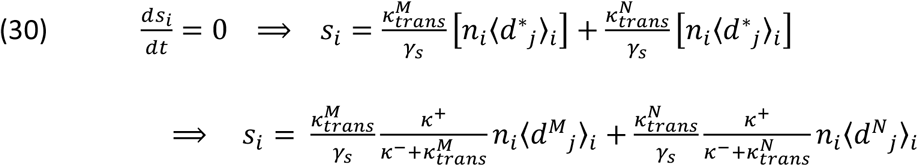

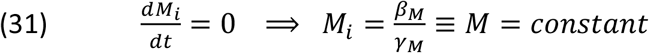
3. We define:

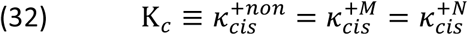

as CI rate,

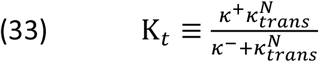

as Trans-activation rate,

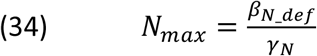

as maximum Neur level at steady state, Where *β_N_def_* is the Maximum Neur expression rate from the default parameters set, such that *β_N_* = *β_N_def_* by default. We use *β_N_def_* as a parameter in the next definition (equation (35)), and distinguished it from the more general *β_N_* (in equation (17));

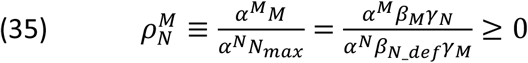

is the ratio between Mib1 and Neur-mediated Delta ubiquitylation,

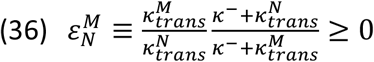

is the ratio between trans-activation from Mib1- to Neur- ubiquitylated Delta.

The equations after simplification become:

##### Ligands and receptor levels

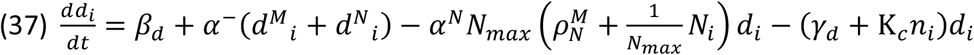

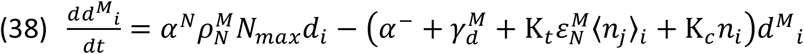

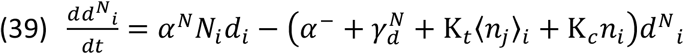

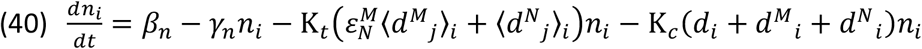

##### Intracellular components levels

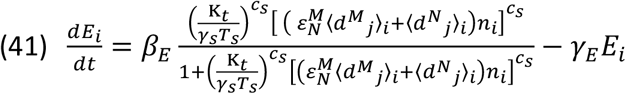

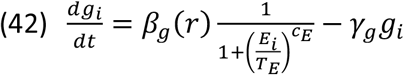

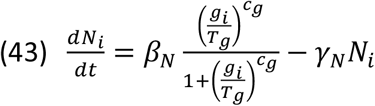

#### Non-dimensionalization

We further reduce the number of parameters in equations (37) to (43) by rescaling the variables using the next assumptions and transformations.

We assume a spatially uniform initial Notch levels, *n*_0_, that is equal to the threshold for promoting *E*(*spl*) expression:

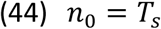

For simplicity, we assume that the degradation rates are equal among all of the variables:

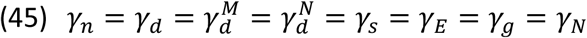

Note that otherwise additional ratio factors need to be introduced.

We normalize all of the levels by the initial Notch level, *n*_0_, and scale the time by the E(Spl) degradation rate, *γ_E_* (which is assumed equal to all other degradation rates). Therefore, the following transformations (scaling):

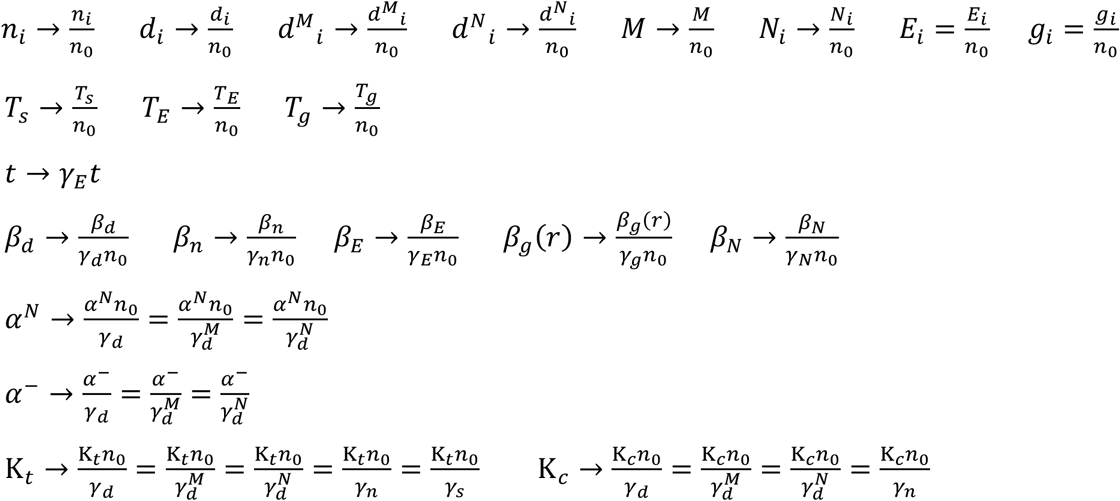

The dimensionless equations after simplifications are:

##### Ligands and receptor levels

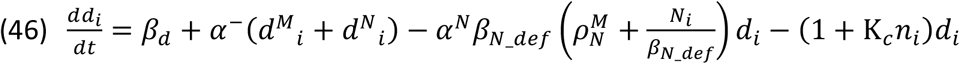

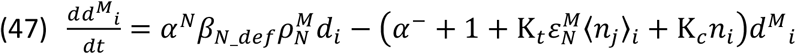

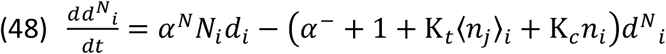

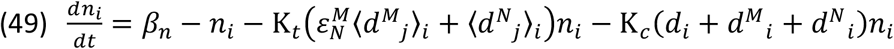

##### Intracellular components levels

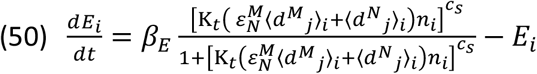

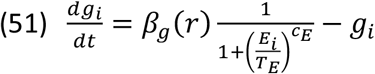

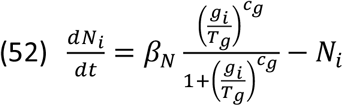

#### Steady state analysis (outside the PNC)

To further reduce the number of free parameters, we analytically solve the dimensionless equations, (46) to (52), at steady state and gain the relations between parameters (Figure 1S1A). Since the notum consists of areas with a very low ac/sc expression rate and levels, outside the PNC, we assume a spatial steady state in this area where all of the variables are at a constant level:

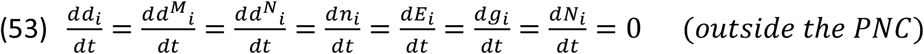

Based on observations that Dl levels are not regulated by ac/sc [15] we assume for simplicity a uniform spatial distribution of Dl and Notch. Therefore:

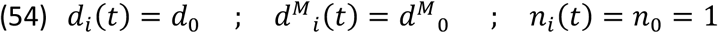

Where *d*_0_, 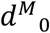 and *n*_0_ are the initial levels of Dl, Mib1 mediated ubiquitylated Dl and Notch, respectively (uniformly expressed among all of the cells).

We also assume that the ac/sc expression rate is equal for all of the cells in this area.

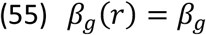

Neur is not expressed due to the very low proneural genes expression rate, which we also interpret as ac/sc level being much less than the Neur threshold:

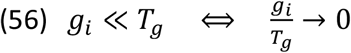

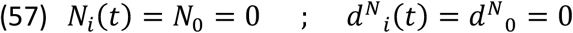

From the assumptions above, (53) to (57), we gain the following relations:

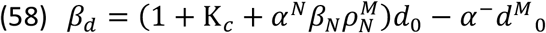

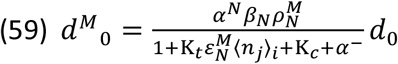

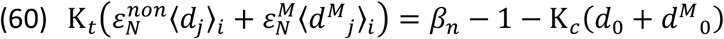

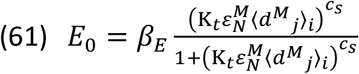

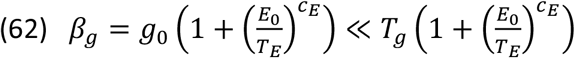

From (58) and (59) we get the condition for the Dl expression rate:

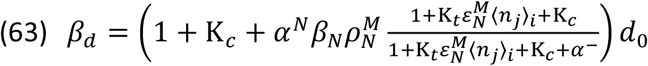

From (58) and (60) we get an expression for the non- and mib1-mediated Notch signal:

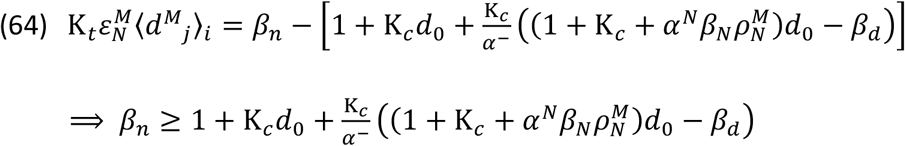

From (61) (63) and (64) we get an expression for the initial E(Spl) level:

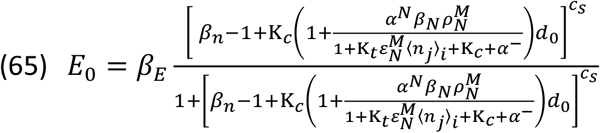

The initial E(Spl) level in (65) can be used for the ac/sc expression rate in (62):

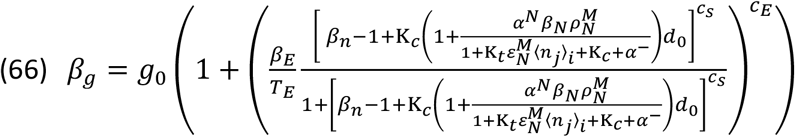

At this point we got the maximum number of relations between the parameters, from the steady state analysis. In the computer simulation part below we will make assumptions for only some of the parameters and then use the relations obtained in this steady state analysis to evaluate the rest of the parameters.

### Computer simulation

The dimensionless equations (46) to (52) are solved numerically, using a standard Matlab ODE solver, and the solution is simulated on a 2D hexagonal lattice consist of 64 cells (Figure 1B). The cells are fixed in size and shape. To avoid edge effect we consider periodic boundaries condition.

As the model contains many unknown parameters, we try to select arbitrary parameters that capture the main feature observed experimentally. Note that some of the parameters could be constrained by experimental observations, while others need to be selected somewhat arbitrarily.

In the next subsections we will determine the set of parameters and initial conditions for the TCS model. All of the parameters and initial conditions used are summarized in table S1.

#### Parameters and initial conditions determination

In this section and the next we complete our assumptions from the steady state analysis above to conclude the parameters and initial conditions set for the TCS model.

We assume a uniform distribution of Notch and Dl along the cell’s boundaries. Since we simulate cells that are arranged in a hexagonal lattice, in the steady state analysis the sums of the Notch, Delta and Mib1 ubiquitylated Dl levels from cells *j* on the boundaries neighbouring cell *i*, are given by:

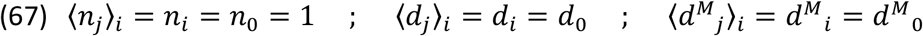

From the relations in (60) and (63), and assumption in (67) we get a relation for the Notch expression rate:

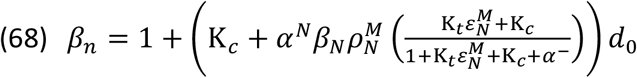

To determine a value for the Neur and E(Spl) expression rates we assume, for simplicity, that they are equal to the Dl expression rate. Using equation (63) and the assumption in (67) we get the following relation:

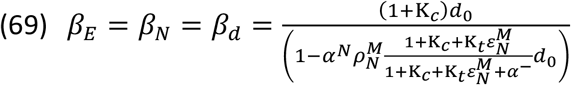

We assume equal initial levels of Dl and Notch:

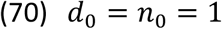

We set the rates for ubiquitylation and trans-activation to be roughly half the Dl expression rate:

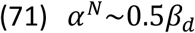

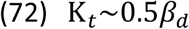

We set the rates for deubiuitylation and CI to be half of the ubiquitylation and transactivation rates:

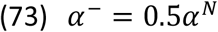

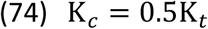

To generate a relatively strong difference between cells that start to express Neur and cells that do not, we set a high Hill coefficient factor for the Neur expression rate:

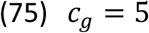

The rest of the Hill coefficients are set to a weaker degree:

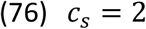

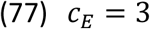

We assume that the initial E(Spl) levels are above the threshold for inhibiting ac/sc expression:

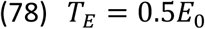

We determine the ac/sc threshold level for promoting Neur expression to be ten times higher than E(Spl) threshold level for inhibiting ac/sc expression:

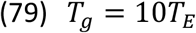

We also set an ac/sc threshold level for SOP selection to be higher than the threshold level for promoting Neur expression:

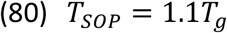

#### Default model assumptions

1. The ratio between Mib1 and Neur ubiquitylation is considered to be between 0 and 1 with priority for Neur ubiquitylation. We therefore set a value of 20% for TCS model, meaning that the probability for a ligand to get ubiquitylated by Neuralized is five times higher than the probability for it to get ubiquitylated by Mib1 (given that Neuralized expression is at its maximum and that Mib1 expression is constant):

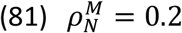
2. Endocytosis rates of ubiquitylated Dl from Mib1 or Neur is assumed to be the same:

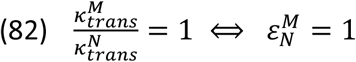

#### Gaussian-like ac/sc expression rate and initial Ac/Sc levels and parameter tuning

Expression rate of ac/sc is not uniform across the notum. For simplicity, and to gain insight on the significance of the parameters, we assume a normalized Gaussian distribution with standard deviation of 1 cell diameter (Note that it is generally not the case in the notum, although some experimentally observed PNC resemble a Gaussian distribution of ac/sc expression; not shown). There is only one cell at the top of the distribution, and the center of this cell is defined as position 0. The distribution of ac/sc expression rate and initial levels drops around the cell at the top with a function:

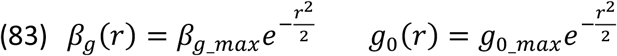

Where *r* is the distance from the cell at the top, given in cell diameter units.

The ac/sc initial levels in the cell at the top of the distribution are set to be in the order of the threshold level for promoting *Neur* expression:

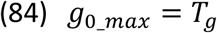

From the steady state analysis above, we get a relation between ac/sc expression rate and initial levels (equation (62)). Yet, this relation is true for a steady state outside the PNC. To allow for increase in ac/sc levels inside the PNC this relation needs to change to an inequality:

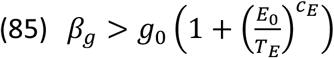

Given the Gaussian-like distribution used in the simulations (see paragraph below), together with the set of parameters obtained above (see also table S1), we tuned the ac/sc expression rate, *β_g_*, inside the PNC to fit the inequality in (85) and to capture the patterns for the wildtype (WT) and mutant phenotypes as observed experimentally (Figure 1B-C, Figure 3B, Figure 5C and Figure 7B). We chose a value that satisfies these conditions:

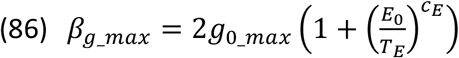

Since we simulate a 2D hexagonal lattice, the distance from the center of one cell to the center of its indirect neighbours (i.e. to its next nearest neighbours, or to its next next nearest neighbours, etc.) is not the same in all directions. We therefore assume a Gaussian-like distribution, where the cell at the top of the distribution has the maximum value of ac/sc expression, *β_g,max_*, and initial levels, *g*_0,*max*_, its nearest neighbours gain the value from the Gaussian in equation (83) where *r* = 1, its next nearest neighbors gain the value where *r* = 2 and all of the other cells have their initial levels of ac/ac and expression rate set to 0.00050 and 0.0046 (respectively) relative to the values in the top cell.

To mimic fluctuations in ac/sc initial levels, we also multiplied *g*_0_ with a noise function which randomizes the value of *g*_0_(*r*) between ±5% of the original value given in equation (83) (and as described above).

#### Simulation Code and summary of parameters and initial conditions

We used custom code in Matlab to run the simulations. The code generating the simulations is given in: https://github.com/Udi-Binshtok/SOP_model_2021 See table S1 for the parameters and initial conditions used in the simulations.

---

**Table S1:**
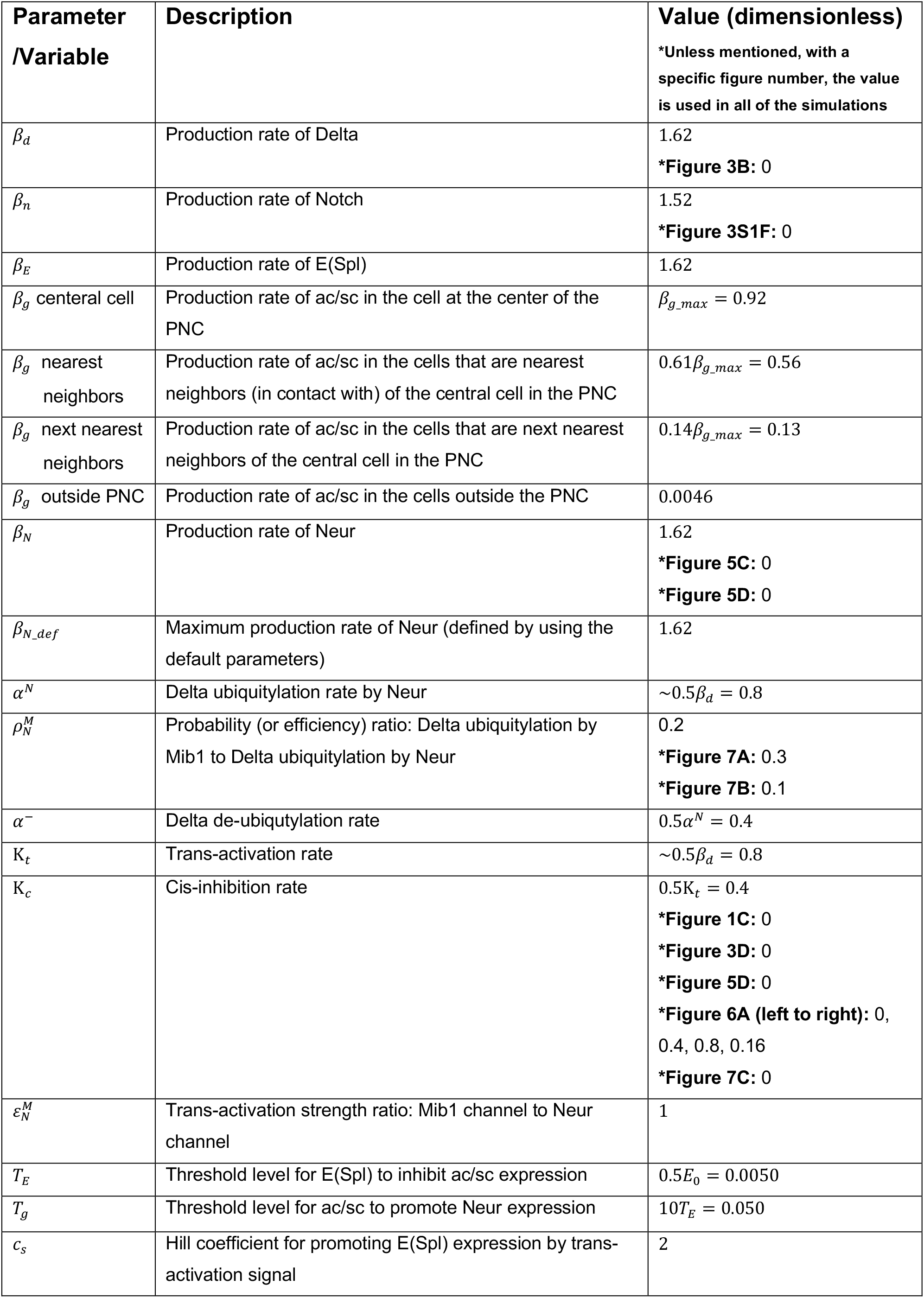
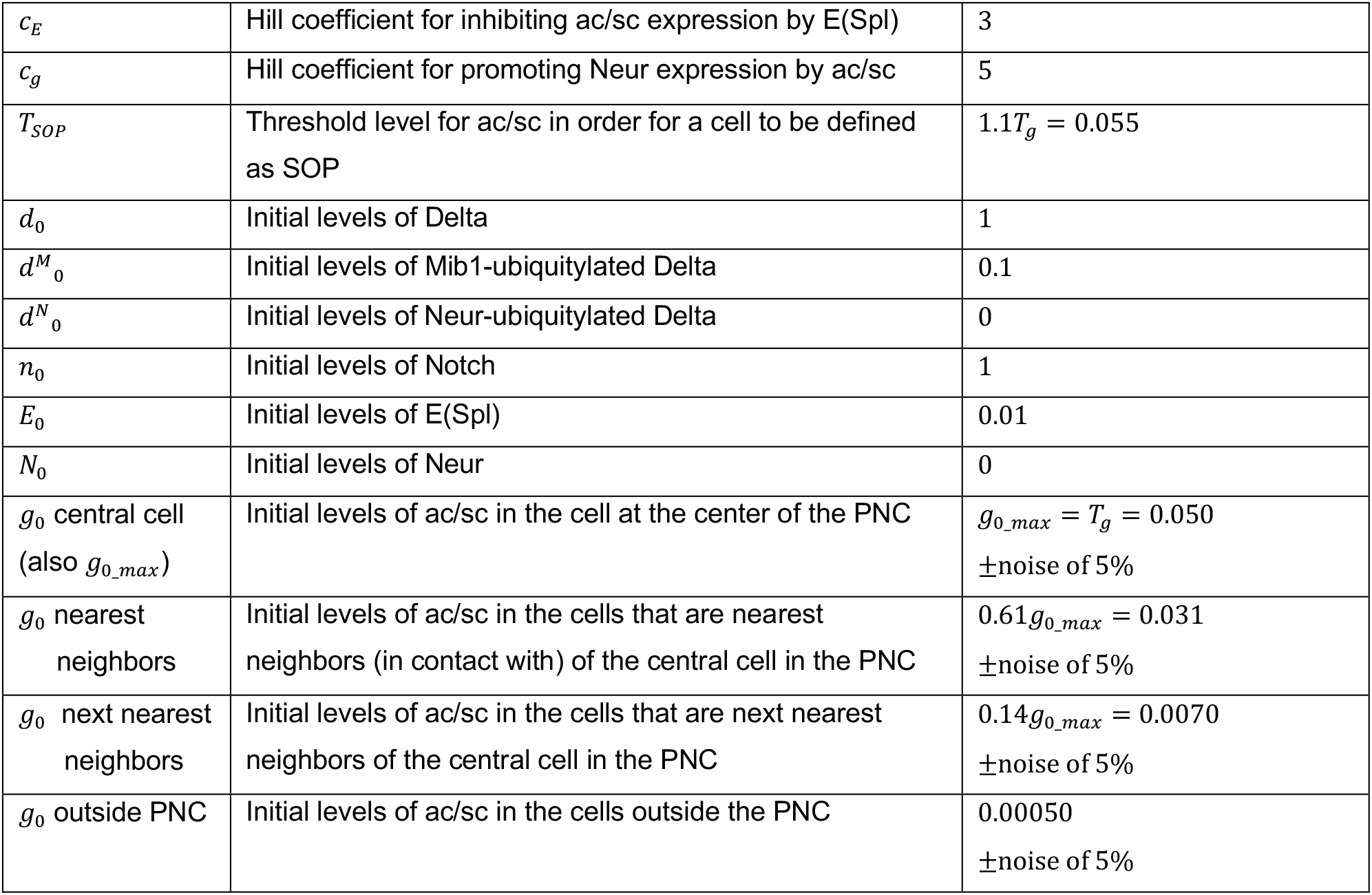
Parameters and initial conditions used in the simulations. These are the parameters used in the final set of dimensionless equations: (46) to (52)

## Supplementary figures

**Figure 1S1:**
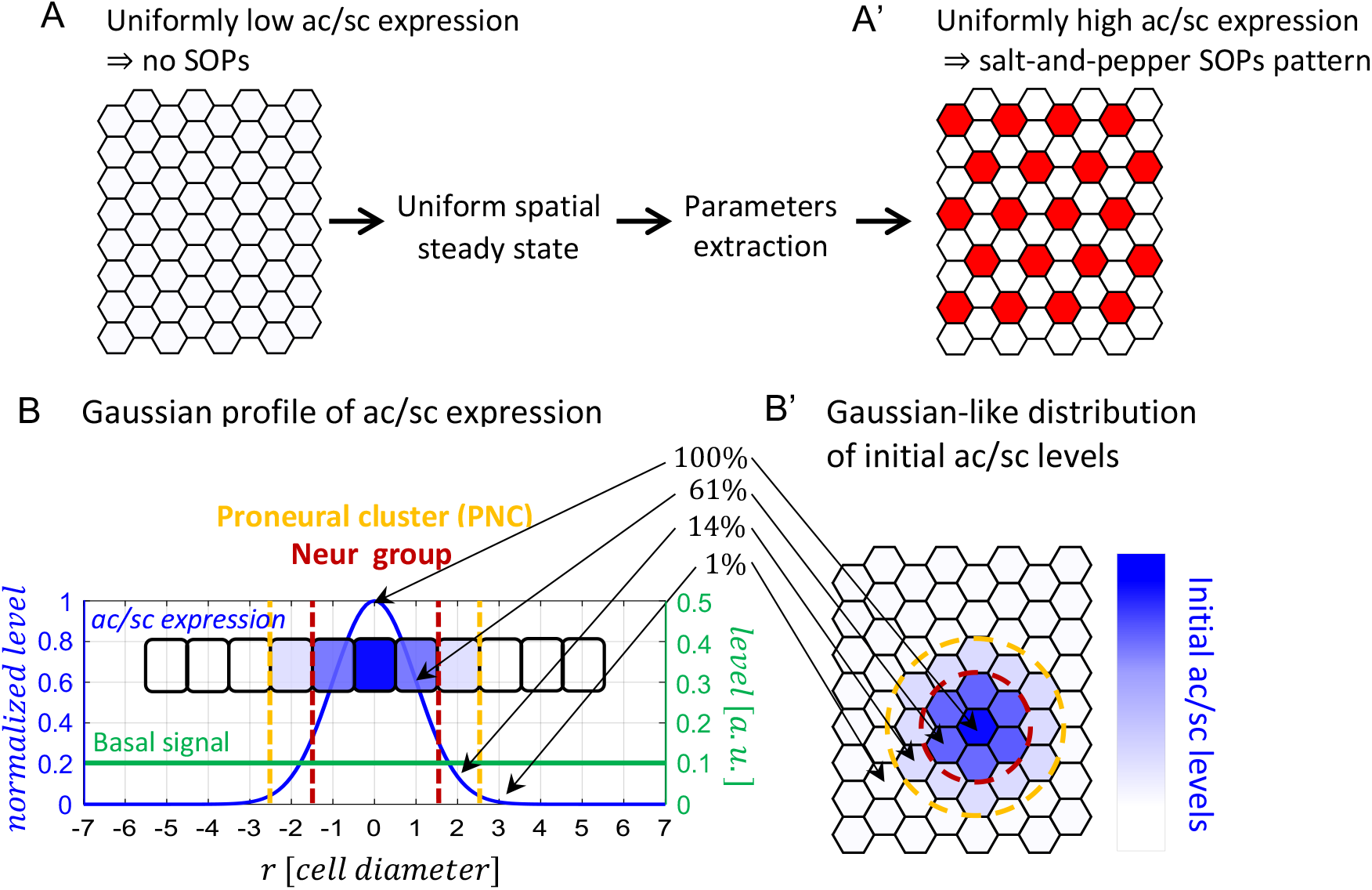
Parameters and initial conditions for the TCS model. **(A-A’)** The model parameters were tuned for domains with either uniformly low (A) or high (A’) ac/sc levels. In (A) ac/sc levels correspond to regions outside the PNC (see parameter “*g*_0_ outside PNC” in table S1) where Neur is not expressed. We first assumed, in this domain, a uniform spatial steady state in the expression of Dl, Notch, E(Spl) and ac/ac (See full analysis in the supplemental material). In (A’) ac/sc levels, across the entire lattice, were set to cross the threshold for Neur expression (see parameter “*g*_0_*max*_” in table S1) to support the formation of a salt-and-pepper pattern (red and white cells in A’). **(B-B’)** Setting the Gaussian profile for ac/sc expression rate and initial levels. The parameters extracted from the steady state analysis in (A) are used in simulations where ac/sc expression exhibit a graded expression profile. The ac/sc production rate is fixed during the simulations and distributed as a normalized Gaussian around one central cell (blue solid line; cells are indicated by rectangles - side view. See also parameter *β_g_* in table S1). A normalized Gaussian profile, 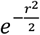, was chosen with a standard deviation of 1 cell diameter. The ac/sc production rate in the central cell (at the pick of the distribution) is set to 100%, while in its nearest neighbors it is reduced to 61%, and in its next nearest neighbors it is reduced to about 14%. In farther away cells the expression rate is reduced to 1 %. Accordingly, the PNC is defined as the central cell together with its nearest and next nearest neighbours (yellow dashed line). Mib1 mediated basal signal is assumed uniform among all of the cells (green solid line). Together, the ac/sc profile and the basal signal define the Neur group inside the PNC (red dashed line). In the WT condition, the Neur group consist of the central cell with only its nearest neighbours. The level of the basal signal is calculated by taking the average (over the total cells population) of the multiplication between the levels of Mib1-ubiquitylated Dl and the levels of Notch - this value is calculated to be 0.1 (See supplemental material and table S1). Top view (B’) shows the initial levels of ac/sc, which are distributed as a Gaussian profile. The PNC and the Neur group are the cells within the yellow and red dashed circles, respectively.

**Figure 2S1:**
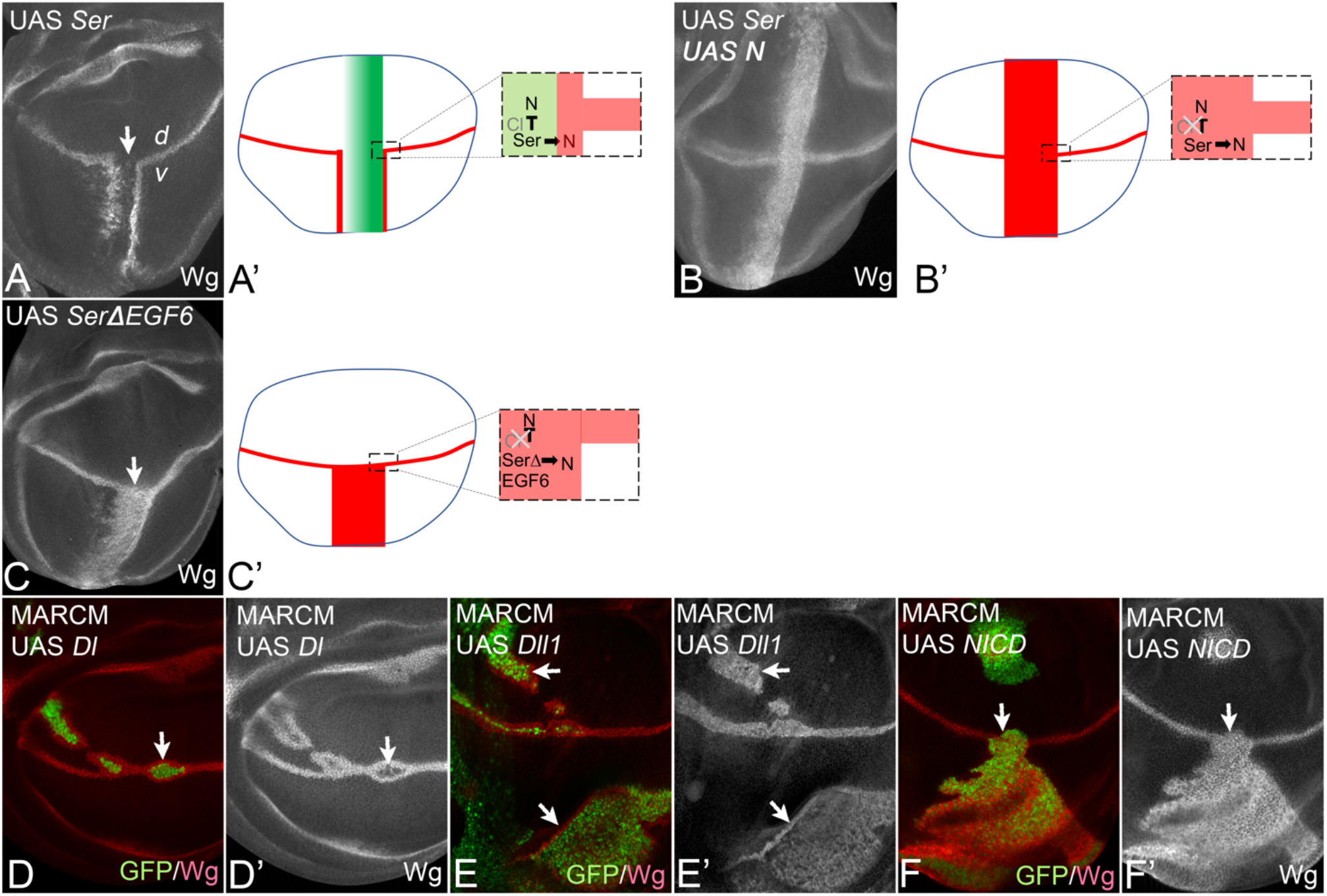
CI by Ser in MARCM clones in the wing. **(A, A’)** Over-expression of UAS *Ser* along the *ptc*Gal4 domain results in the interruption of the endogenous expression of Wg due to CI (arrow). In addition, similar to Dl (Fig. 1B-B’), it induces two ectopic stripes of Wg. For Ser, these stripes are restricted to the ventral compartment. **(B, B’)** The CI of Ser is also suppressed by co-expression of Notch. **(C, C’)** A Ser variant that lacks EGF6, SerΔEGF6, in the extracellular domain does not cis-inhibit (arrow) in a similar manner as Dll1. Only a broad stripe of ectopic expression throughout the *ptc*Gal4 domain is induced, restricted to the ventral compartment. **(D-F’)** MARCM clones expressing Dl (D, D’), Dll1 (E, E’) and NIDC (F, F’). The expression of Dl results in only very weak expression of Wg within the clone, but strong expression in cells adjacent to the clone (arrow), indicating that CI occurs within the clone. In contrast, Dll1 induce high expression of Wg also within the clone showing that Dll1 does not CI also in MARCM clones (arrows in E,E’). A similar behaviour is observed when NICD is expressed with the clone (arrow in F, F’).

**Figure 3S1:**
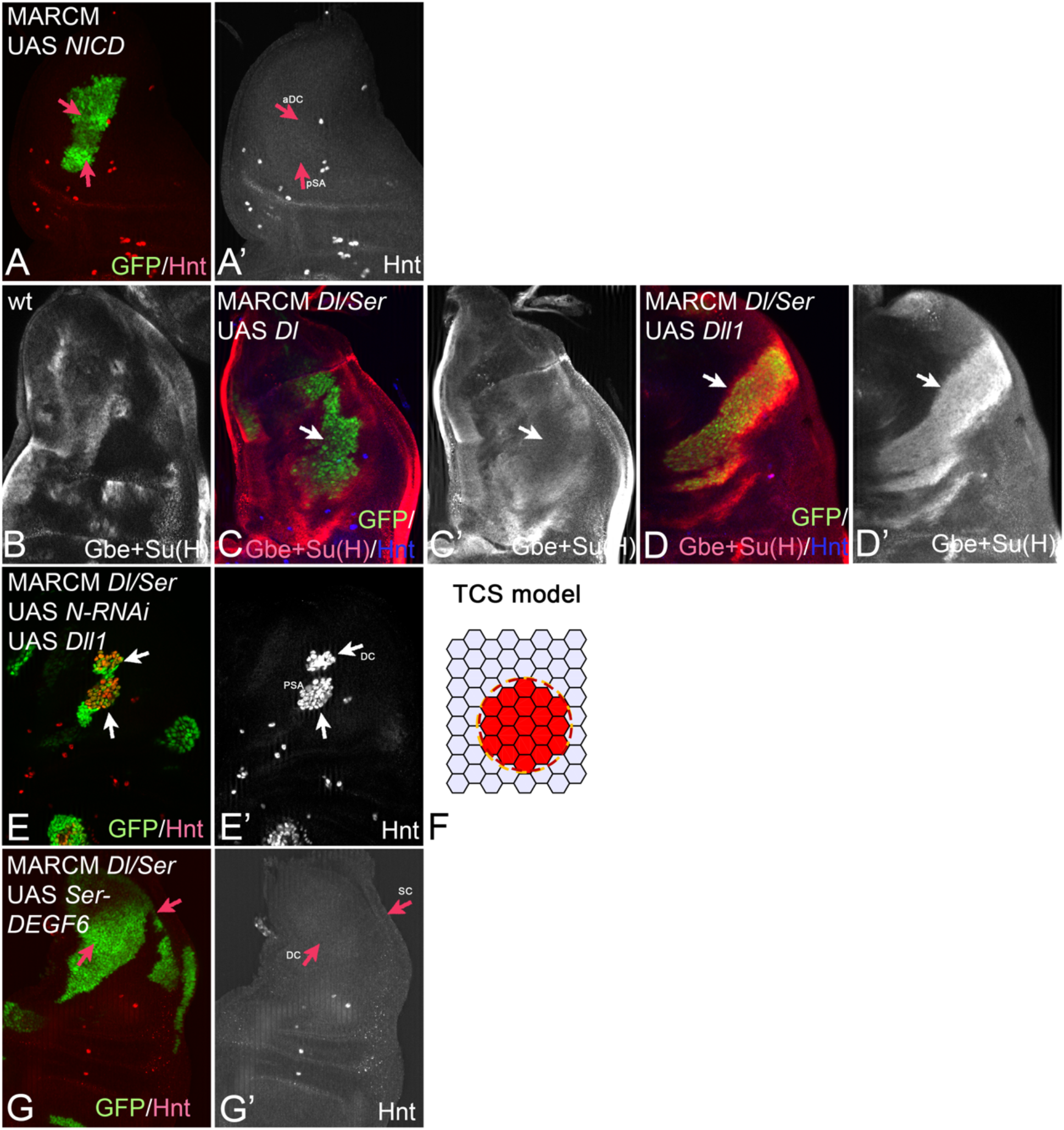
CI is required for the selection of the SOP in the notum. **(A-A’)** MARCM clones expressing NICD. No SOP formation can be observed in the clone cells. (arrows, compare with Fig. 2S1F). **(B-F)** Expression of Dll1 induces higher Notch activity (in trans) than Dl. **(B)** Expression of the Notch activity reporter Gbe+Su(H) in a wildtype notum. **(C, C’)** A wing disc bearing a Dl/Ser MARCM clone that expresses Dl (arrow). No significant elevation of expression of Gbe+Su(H) is observed. **(D, D’)** In contrast, a strong elevation of Gbe+Su(H) expression is observed in clones that express Dll1 (arrow). **(E-F)** The suppression of SOP formation in *Dl/Ser* MARCM clones expressing Dll1 is abolished upon co-expression of N-RNAi. Instead, an excess of SOP formation is observed (neurogenic phenotype, arrows in E, E’). **(F)** An excess of SOP formation is predicted by the TCS model when Notch production rate is set to zero. **(G, G’)** Similar to Dll1 (Fig. 3D-D’”), expression of the non-cis-inhibitory SerΔEGF6 in *Dl/Ser* MARCM clones suppresses SOP formation (arrows).

**Figure 3S2:**
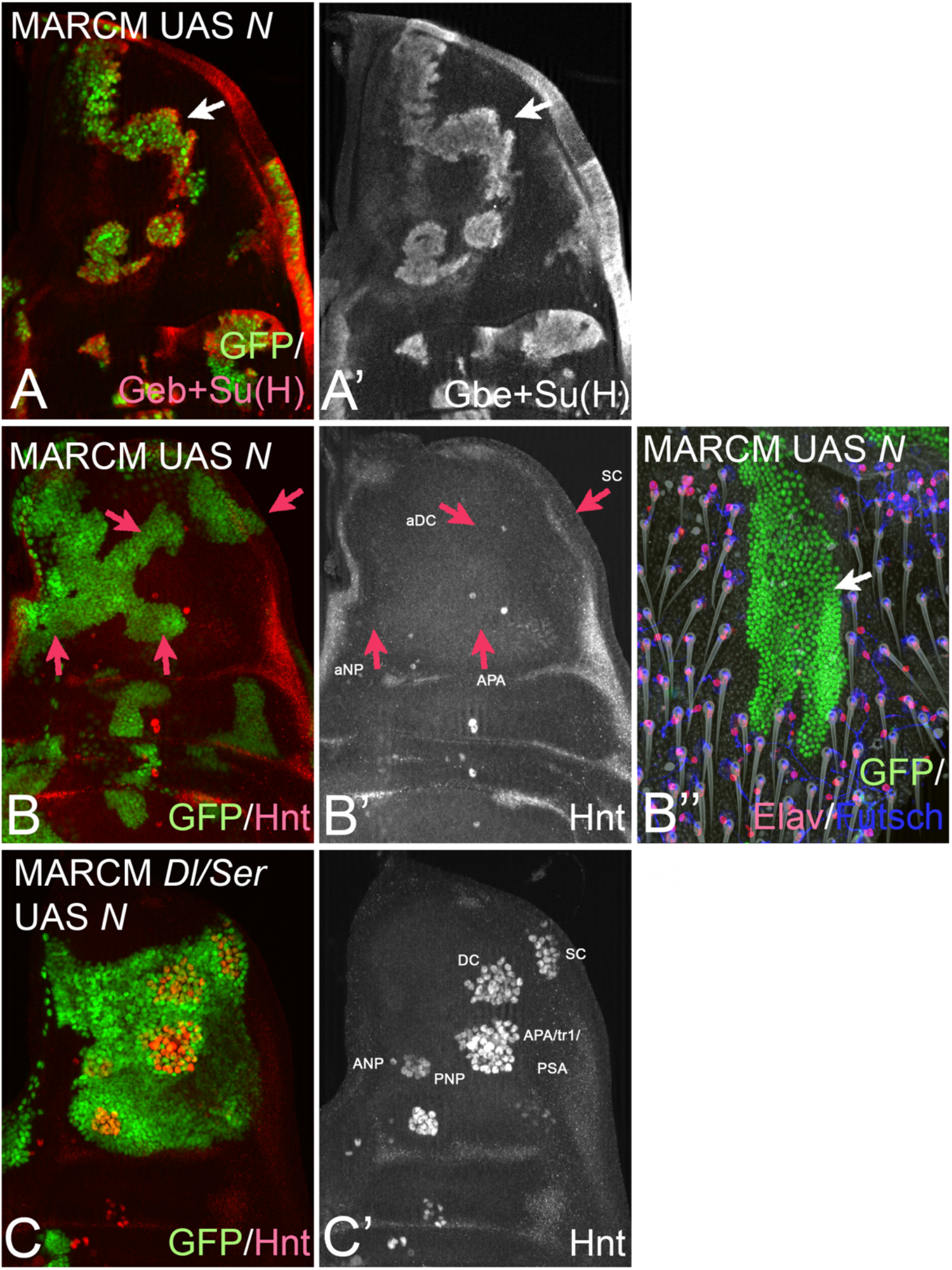
Overexpressing Notch suppresses CI of endogenous ligands. **(A, A’)** Expression of Notch in MARCM clones results in a strong increase of Notch pathway activity, revealed by the induction of high level of expression of Gbe+Su(H) reporter (arrow). **(B-B”)** The expression of Notch suppresses SOP formation in the disc (arrows). Consequently, no bristle formation can be observed in clonal regions in adults (B”, arrow). The additional staining with antibodies against the neural proteins Elav and Futsch reveal that also the internal elements of the bristle organ are absent. The suppression of SOP formation by elevation of Notch levels is abolished if the MARCM clones are additionally mutant for Dl and Ser, indicating the requirement of the ligands for suppression. (C, C’) A large MARCM clone which expresses UAS N and is mutant for *Dl* and *Ser*. A strong neurogenic phenotype can be observed.

**Figure 4S1:**
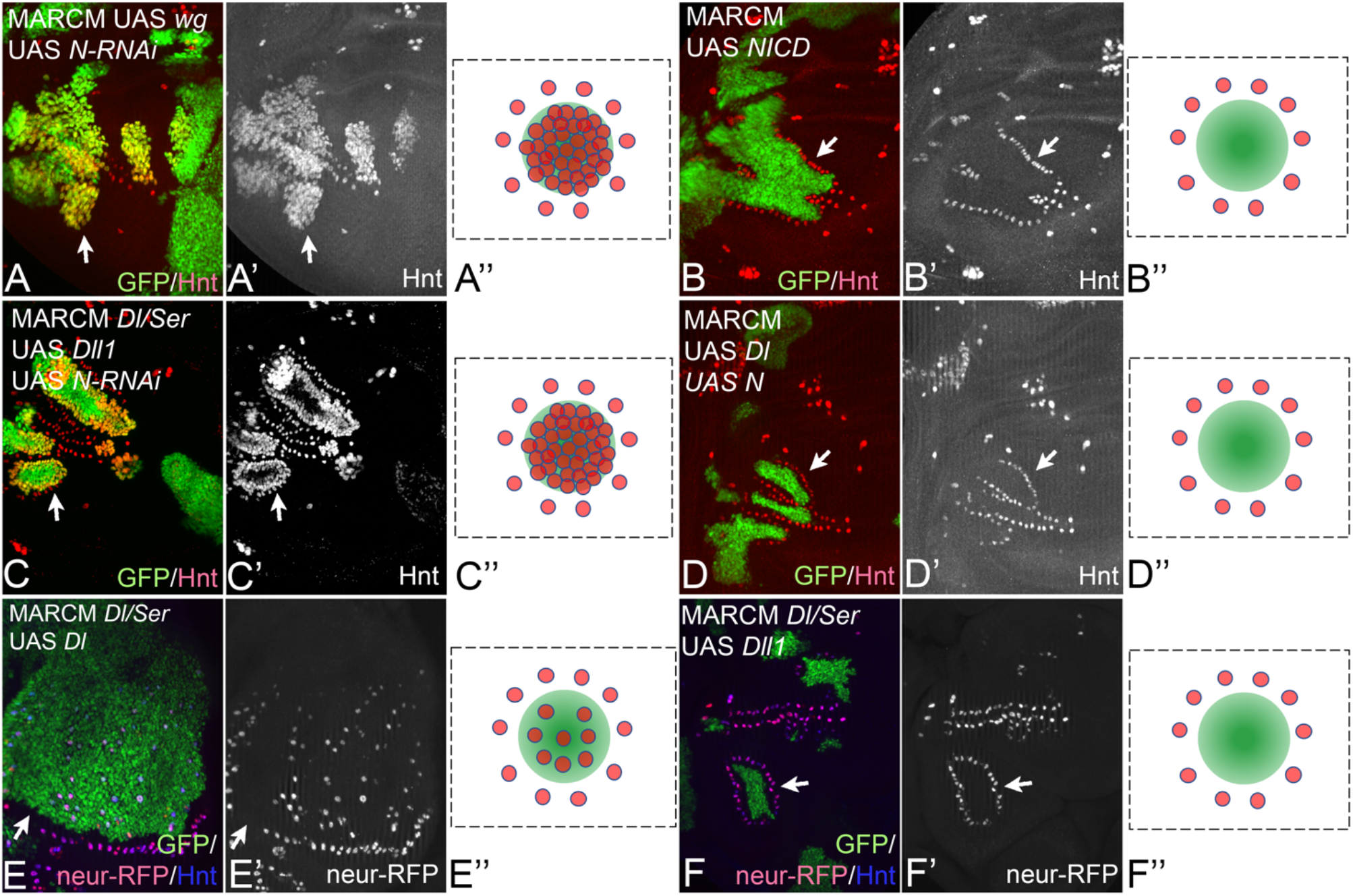
Controls for MARCM clone experiments. **(A-A”)** Wg expressing MARCM clones depleted of Notch display a neurogenic phenotype, indicating that the selection of the SOP seen in the Wg expressing clones depends on Notch signalling (arrow, compare with Fig. 4B-B”). **(B-B”)** Similar to Dll1, expression of NICD in MARCM induces SOP formation only in cells adjacent to the clone (arrow). **(C-C”)** Depletion of Notch in *Dl/Ser* MARCM clones that express Dll1 abolishes the suppression of SOP formation. Instead, a neurogenic phenotype is observed, indicating that Dll1 suppresses SOP formation within the clone via the activation of the Notch pathway. **(D-D”)** Co-expression of Dl and N in MARCM clones causes a phenotype similar to that of expression of Dll1. SOP formation is observed in cells adjacent to the clone. In the clone, SOP formation is abolished because of the suppression of CI of Dl by high levels of Notch. **(E-E”)** The ectopic SOPs within *Dl/Ser* MARCM clones that express Dl also express Neur, suggesting the formation of the Neur-group. **(F-F”)** This is not observed when Dll1 is expressed within *Dl/Ser* MARCM clones, suggesting that the high activity of Notch induced by Dll1 prevents the formation of the Neur-group.

**Figure 5S1:**
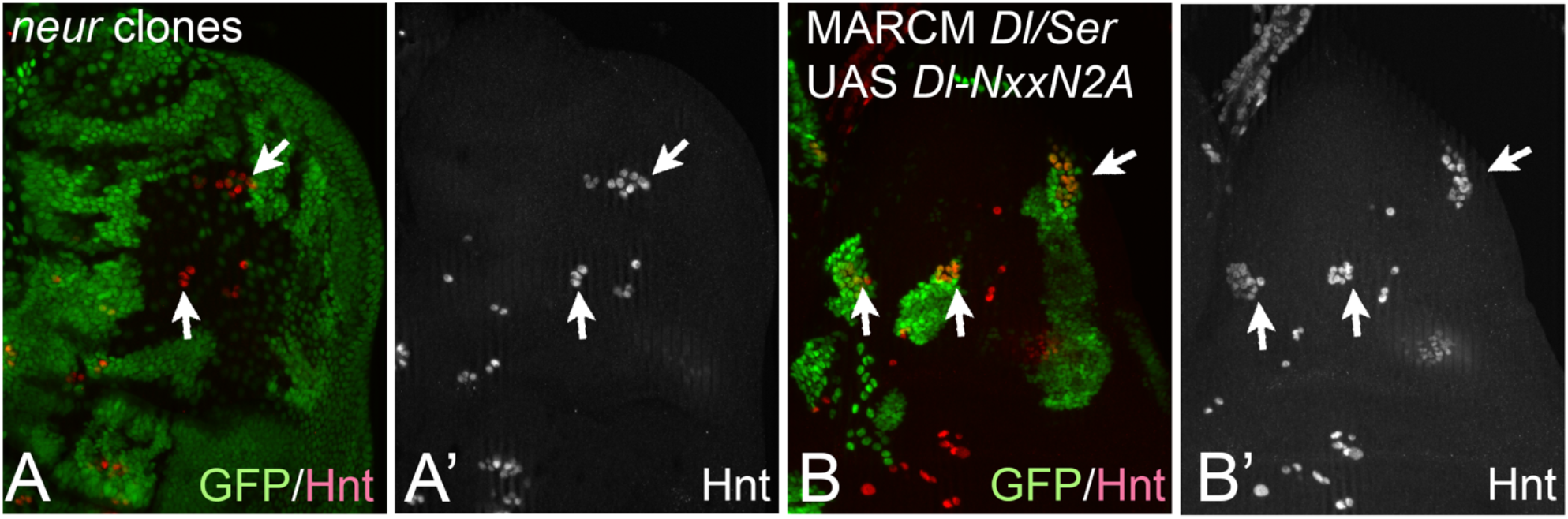
(A-A”) The phenotype of *neur* mutant clones. The clones are labelled by absence of GFP. A weak neurogenic phenotype is observed (arrows). In contrast to Dl, expression of a Dl variant without Neur binding site, Dl-NxxN2A, cannot rescue SOP selection in *Dl/Ser* mutant MARCM clones. As in the case of *neur* clones, a weak neurogenic phenotype is observed (arrows).

## Notes

### Competing Interest Statement

The authors have declared no competing interest.

